# CTx001 for geographic atrophy: a gene therapy expressing soluble, truncated complement receptor 1 (mini-CR1)

**DOI:** 10.1101/2025.06.03.657557

**Authors:** Sonika Rathi, Athanasios Didangelos, Sofiya Pisarenka, Rachel Green, Peter Emery-Billcliff, Nakul Patel, Pamela Whalley, Parisa Zamira, Viranga Tilakaratna, Ewa Szula, Rafiq Hasan, Mustafa Munye, Richard D. Unwin, Paul N Bishop, Dennis Keefe, Simon J Clark

## Abstract

Over-activation of the complement system is strongly associated with geographic atrophy (GA), a late-stage form of age-related macular degeneration (AMD) and major cause of blindness. Here we report the development of a candidate GA treatment that is a potent complement modifier (mini-CR1) that addresses consequences of complement over-activation including membrane attack complex (MAC) formation. Mini-CR1 acts a potent cofactor for factor I driven proteolysis of C3b leading to iC3b and then C3dg formation; iC3b itself remains a potent opsonin. As well as inhibiting the alternative complement pathway, mini-CR1 acts as a cofactor for C4b degradation thereby inhibiting the classical pathway. Mini-CR1 prevents MAC formation in activated human serum with an IC_50_ of 125 nM. Mini-CR1 was shown to cross human Bruch’s membrane *ex vivo*, implying the ability to cross into the choroidal space. Transduction of RPE cell lines with rAAV2-mini-CR1 (CTx001) resulted in dose-dependent transcription, and both basolateral and apical secretion of mini-CR1 by monolayers of human iPSC derived RPE cells. Mini-CR1 transduction of ARPE19 cells resulted in increased consumption of C3b and iC3b1 in activated culture media and decreased MAC formation on the cell surfaces. Subretinal injection of CTx001 in rats resulted in dose-dependent mini-CR1 production as demonstrated by solid-phase immunoassay. MAC formation following laser induced CNV in a rat model was reduced by 75% in CTx001-treated animals relative to null vector (*p*<0.01). This first generation of CTx001 represents a potent single administration complement modifier capable of effectively addressing pathologic complement amplification in the retina/choroid.

## Introduction

Age-related macular degeneration (AMD) is a degenerative blinding disease that manifests itself through the progressive destruction of the macula, the central region of the retina responsible for high-resolution vision. The condition, which constitutes the leading cause of blindness in developed countries ^1,2^, has two late-stage forms: wet AMD, characterised by the formation of choroidal neovascularisation (CNV) and geographic atrophy (GA; also called late dry AMD), in which patches of retinal cell death form. Since the introduction of anti-VEGF therapies as treatments for CNV, the wet form of AMD has been clinically well-managed ^3^. In 2023 two treatments for GA that target the complement system called pegcetacoplan and avacincaptad pegol ^4^ were approved by the US FDA. These therapies that are delivered by regular intravitreal injection inhibit complement and have been shown to slow the progression of GA lesions.

Evidence for the involvement of the complement system in AMD has emerged over nearly two decades of genetic, biochemical and human tissue studies. The retina (neurosensory retina and retinal pigment epithelium (RPE)) along with the underlying Bruch’s membrane and choriocapillaris (the outer retinal blood supply) are where tissue damage occurs in AMD and there is evidence for complement mediated damage in all these layers (Figure 1A). However, recent evidence suggests that complement overactivation occurs both in the neurosensory retina and the choriocapillaris layer, particularly the extracellular matrix (ECM) surrounding this layer of capillaries and its associated Bruch’s membrane ^5–8^.

**Figure 1.**
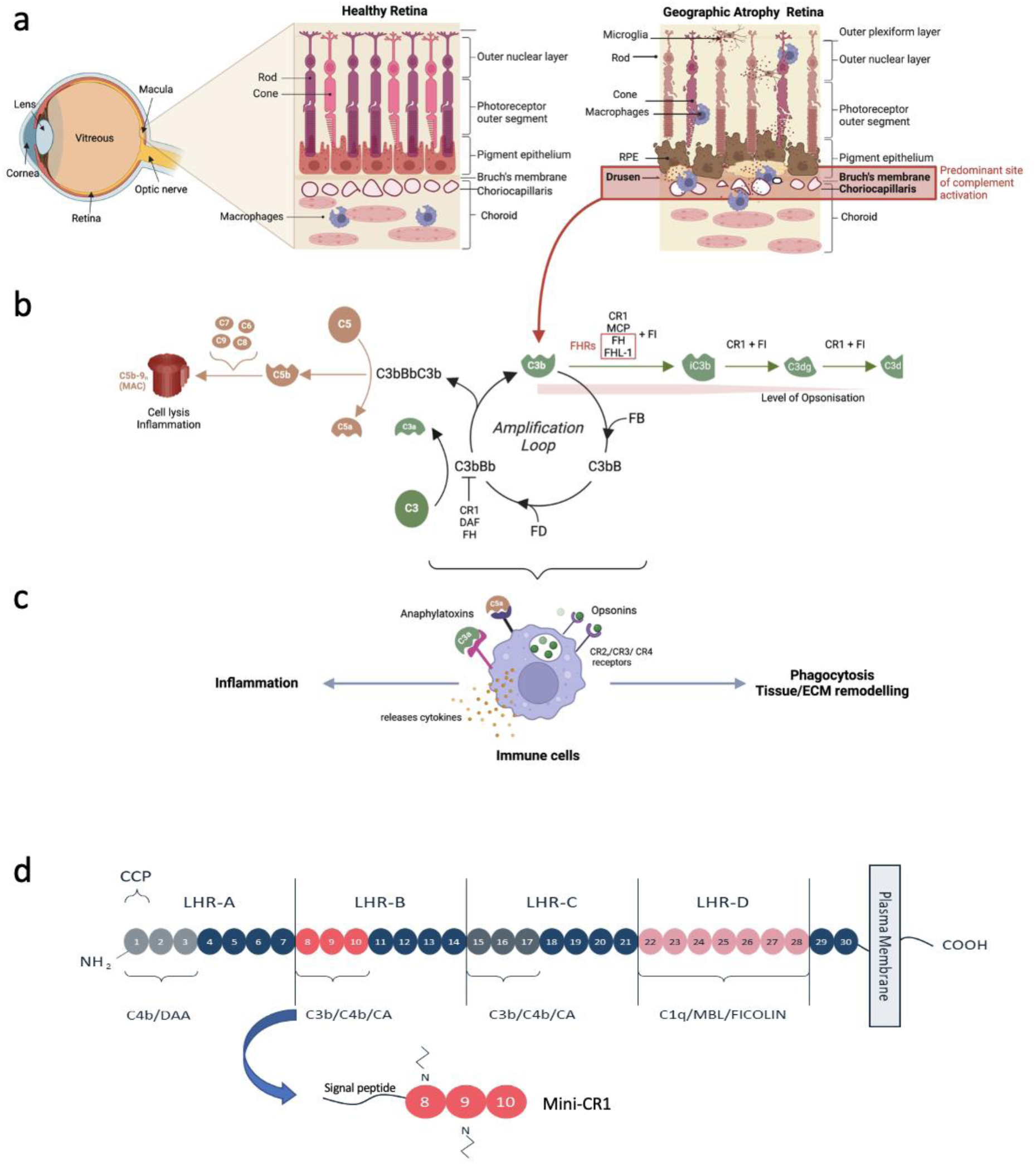
Schematic representation of the human eye, with a focus on retinal changes in geographic atrophy (GA) and complement activation. a) The human retina in the posterior part of the eye consists of the light-sensitive neurosensory retina and RPE, underlying the RPE is Bruch’s membrane and the inner layer of the choroid, called the choriocapillaris. GA occurs in the central part of the retina called the macula. During the formation of GA lesions, which often form in areas with extracellular deposits in Bruch’s membrane called drusen, there is atrophy of the RPE, photoreceptors and the choriocapillaris. This micro-environment is characterized by the presence of activated complement products in and around the extracellular matrix (ECM) of Bruch’s membrane and the choriocapillaris that promotes both inflammation and the ingression of cells including macrophages and microglia. b) Complement can be activated through three separate pathways ^46^, but the central protagonist remains the deposition of C3b onto cells and ECM. The binding of factor B (FB) and processing by factor D (FD) creates a C3 convertase (C3bBb), which enzymatically cleaves circulating C3 into C3b, feeding into the amplification loop. Subsequently, a C5 convertase is also formed (C3bBbC3b) which cleaves circulating C5 into C5b and triggers the formation of the membrane attack complex (MAC, or C5b-9_n_) that causes cell lysis and contributes to a pro-inflammatory environment. Processing of C3 and C5 also produces C3a and C5a: two potent anaphylatoxins, which act as chemoattractants that recruit circulating immune cells to the site of complement activation. Complement activation is regulated by the protease Factor I (FI) and this requires several one of several cofactors including Factor H (FH), Factor H-like protein 1 (FHL-1), Membrane co-factor protein (MCP), and Complement receptor 1 (CR1). Increasing concentrations of FHR proteins can interfere with the co-factor activity of FH and FHL-1 (red box). Together with a cofactor, FI cleaves C3b into inactive C3b (iC3b) which can no longer contribute to the amplification loop of complement activation. Both C3b and iC3b are potent opsonins: molecules that label a surface for phagocytosis by the immune cells recruited by the C3a/C5a anaphylatoxins. Opsonisation is finally switched off through further degradation of iC3b by FI into C3dg and eventually C3d: CR1 is the only FI cofactor able to support this step. c) Engagement of the C3a or C5a receptors on recruited immune cells stimulates degranulation and release of proteases and pro-inflammatory cytokines. Opsonin engagement with specific receptors initiates phagocytosis and tissue/ECM remodelling. d) Native CR1 is a transmembrane glycoprotein comprised of 30 complement control protein (CPP) domains organized across four long homologous repeats (LHRs). Mini-CR1 comprises CCPs 8-10 (depicted in red), which contain the C3b/C4b binding domain, the pre-protein form has an N terminal signal peptide to enable secretion from cells. DAA = Decay accelerating activity; CA = cofactor activity; N = N linked glycosylation sites.

The complement system consists of the classical, lectin and alternative pathways that all feed into a central C3 amplification loop. The C3 amplification loop feeds into the C5 amplification loop which in turn activates the terminal pathway and membrane attack complex (MAC) formation (Figure 1B). In the C3 amplification loop, C3 is broken down into C3a and C3b. C3b binds to factor B to form a convertase which in turn stimulates more C3b production and downstream activation of the complement pathway *via* C5 cleavage, resulting in C5a production and MAC (C5b9n) formation. The anaphylotoxins C3a and C5a are potent mediators of inflammation and activators of immune cells (Figure 1C). Pegcetacoplan and avacincaptad pegol act by inhibiting the breakdown of C3 into C3a and C3b, and C5 into C5a and C5b respectively.

The C3 amplification loop is downregulated by the breakdown of C3b by the enzyme complement factor I (FI) into iC3b, a process that requires the presence of a FI cofactor. Cofactors include the soluble proteins factor H (FH) and its truncated variant factor H-like protein (FHL-1), and the cell membrane-bound complement receptor 1 (CR1). FH/FHL-1 allow the breakdown of C3b as far as iC3b, but this breakdown product remains a potent opsonin. In contrast, CR1 permits the further FI-mediated degradation of iC3b into C3c/C3dg towards lesser opsonin potency, a reduction in pro-inflammatory stimuli, and greater inactivation. Variations in the *CFH* gene that result in decreased function of the FH and FHL-1 proteins represent a major genetic risk factor for AMD. In addition, there are 5 FHR proteins (FHR1-5) that inhibit the actions of FH and FHL-1 thereby impeding C3b breakdown and driving complement activation; variations in and around genes encoding the FHR proteins, including *CFHR1*, *CFHR2*, *CFHR4* and *CFHR5* have been shown to be associated with, and indeed causal in AMD ^9–13^.

To develop a treatment for GA we elected to use the approach of enhancing cofactor activity, thereby generally suppressing complement activation and specifically restoring deficient cofactor activity in patients. We based our approach on CR1, as it has potent cofactor activity and is the only natural complement regulator capable of degrading iC3b. However, the native CR1 protein is a large (∼220kDa) glycoprotein comprising thirty complement control protein (CCP) domains (Figure 1D), whereas we aimed to use a small, secreted protein that can traverse Bruch’s membrane and thereby treat complement overactivation on both the retinal and choriocapillaris sides of Bruch’s membrane. Therefore, we elected to use one of the two elements within CR1 that provide its cofactor activity i.e. CCPs 8-10. Delivery of this protein using subretinal gene therapy has the advantage of being a potential one-off treatment rather than the frequent intravitreal injections required for current therapies. In addition, production and secretion of the protein from RPE cells could help avoid the blood-retina-barrier and support effective distribution to both the retinal and choroidal compartments. To allow the protein to be secreted, the FH signal peptide was added, and the resultant therapeutic protein was called mini-CR1 (Figure 1D). An rAAV2 self-complimentary vector was developed that encoded mini-CR1 (CTx001), and below the biological activity and therapeutic potential of this vector and mini-CR1 are investigated.

## Materials and methods

### Purified recombinant mini-CR1 protein production in HEK293T cells

To produce purified, recombinant mini-CR1 protein, cDNA encoding CCPs8-10 of CR1, which includes a TEV-cleavable N-terminal 6xHis tag, was incorporated into a pcDNA3.1 vector and transfected into HEK293T cells using previously described expression protocols ^14^. Further details on the expression and purification of recombinant mini-CR1 are provided in the Supplementary Methods.

### C3b/C4b degradation assays

*In vitro* digestions of human C3b and C4b were conducted to assess the cofactor activity of mini-CR1 in the presence of human FI (CompTech, USA) and the results were analysed by densitometry of chemiluminescence signals following Western blotting; see Supplemental Methods for further details.

### Alternative complement pathway inhibition assays

The Wieslab Alternative Complement Pathway Assay (SVAR COMPLAP330, Sweden) was employed to assess the mini-CR1 complement inhibitory activity in human and Cynomolgus Monkey serum and the Hycult Alternative Complement Pathway Assay (Hycult Biotech, The Netherlands) was employed for murine serum analysis. See supplemental methods for further details.

### Binding affinity studies using Biolayer Interferometry

Measurements of binding affinities for mini-CR1, FHL-1, and FH to C3b were performed using an Octet RH384 Biolayer Interferometry (BLI) system (Satorius, Germany). Streptavidin biosensors were prepared by immobilizing human C3b (Merck, UK) that has been biotinylated using an EZ-Link Sulfo-NHS Biotin kit (ThermoScientific, UK) according to the manufacturer’s instructions, as described previously for other proteins ^15^. Then mini-CR1, FHL-1 and FH were separately passed over the C3b coated sensors to allow calculation of association and dissociation rate constants. For further details see Supplementary Methods.

### Ussing Chamber Diffusion

The diffusion properties of mini-CR1 across Bruch’s membrane were characterized *ex vivo* in Ussing chambers (Harvard Apparatus, USA) with enriched human Bruch’s membrane as described previously ^16^. For further details see Supplementary Methods.

### rAAV production and transduction of RPE cell lines

rAAV constructs were prepared by SignaGen Laboratories using their proprietary rAAV Production Service. To assess the efficiency of the CTx001 rAAV constructs in expressing mini-CR1, transduction experiments were conducted in ARPE19, hTERT-RPE1 and primary RPE cells. For further details of RPE cell transduction and analysis of expression see Supplementary Methods.

### Human iPSC-RPE culture, transduction and analysis of directionality of mini-CR1 secretion

Human induced pluripotent stem cell (iPSC) derived RPE cells were cultured and then seeded onto Matrigel® coated transwell-12 inserts. They were allowed to form a monolayer and tight junctions were assessed using occludin and ZO-1 immunostaining. Secretion of mini-CR1 into the apical and basolateral facing media was measured using mesoscale [MSD] electrochemiluminescence. See Supplemental Methods for further details.

### *In vivo* subretinal delivery of viral vectors

The *in vivo* rat studies were performed by Powered Research, USA. Animals were anesthetized and placed under a stereoscope (Leica Microsystems) and a drop of iodine was applied on the cornea and allowed to spread evenly (Minims Povidone Iodine 5%, Laboratoire Chauvin S.A.). A sclerotomy in the temporal side was performed with a 30G needle in order to expose the choroid. The cornea was punctured to reduce the intraocular pressure. A microsyringe (Hamilton Bonaduz AG) was filled with rAAV2 particles and introduced into the subretinal space through the exposed choroid. The solution was injected into the subretinal space for 10 sec, and the needle was kept in place for an additional 30 sec before being removed. The success of the injection was confirmed using in vivo SD-OCT imaging (Bioptigen Envisu R2210. Bioptigen Inc./Leica Microsystems). Chloramphenicol ointment was applied after the injection (Oftan Dexa-Chlora, Santen Oy). CNV induction was performed 28 days after administration of rAAV2 particles. Rats were anesthetized, and pupils were dilated by topical instillation of 1% tropicamide (Tropicamidum WZF 1%: Polfa S. A., Poland or Mydriacyl, s.a. Alcon-Couvreur n.v., Belgium). Carbomer gel (Lakripos, UrsaPharm, Germany) was applied to the right eye (OD) and a coverslip was used to applanate the cornea. Three laser lesions were placed around the optic nerve head of the right eye (OD) using a 532 nm diode laser (Lumenis^®^, Novus Spectra) and using the following standardized settings: spot size: 100 µm; power: 100 mW; time: 100 ms. For staining of choroidal flat mounts with fluorescein-labeled isolection B4 see Supplementary Methods.

## Results

### Mini-CR1 drives proteolysis of C3b, iC3b and C4b by FI

Recombinant mini-CR1 is efficiently secreted by mammalian cells, and once purified appears as two bands on reducing SDS-PAGE gels (Supplementary Figure 1). This dual band appearance is due to the presence of two N-glycosylation sites in mini-CR1, with the higher molecular weight band having both occupied, whereas the lower band only has one occupied: the two bands resolve to a single band after treatment with PNGase F to remove glycosylation (Supplementary Figure 1).

C3b breakdown assays demonstrated that mini-CR1 at concentrations as low as 62 ng/ml (2.1nM) caused substantial consumption of C3b with the production of iC3b, C3dg and C3c (Figure 2A-C). As the concentration of mini-CR1 increased the consumption of C3b became more complete, as did the breakdown of iC3b to C3dg and C3c. C4b degradation also occurred in a dose-dependent pattern with increasing doses of mini-CR1 resulting in decreased C4b and an increase in C4d fragments, although the digestion efficiency was lower than for C3b (Figure 2A and D). In contrast to mini-CR1 and CR1, other FI cofactors only facilitate the cleavage of C3b into iC3b. This is demonstrated for FHL-1 (Supplementary Figure 2) and known to be the case with FH and CD46/MCP ^17^. Mini-CR1 shares the same capacity as soluble CR1 (a shortened form of CR1 that lacks a cell membrane binding domain found in very low concentration in blood) to drive the breakdown of C3b into C3dg (Supplementary Figure 2).

**Figure 2:**
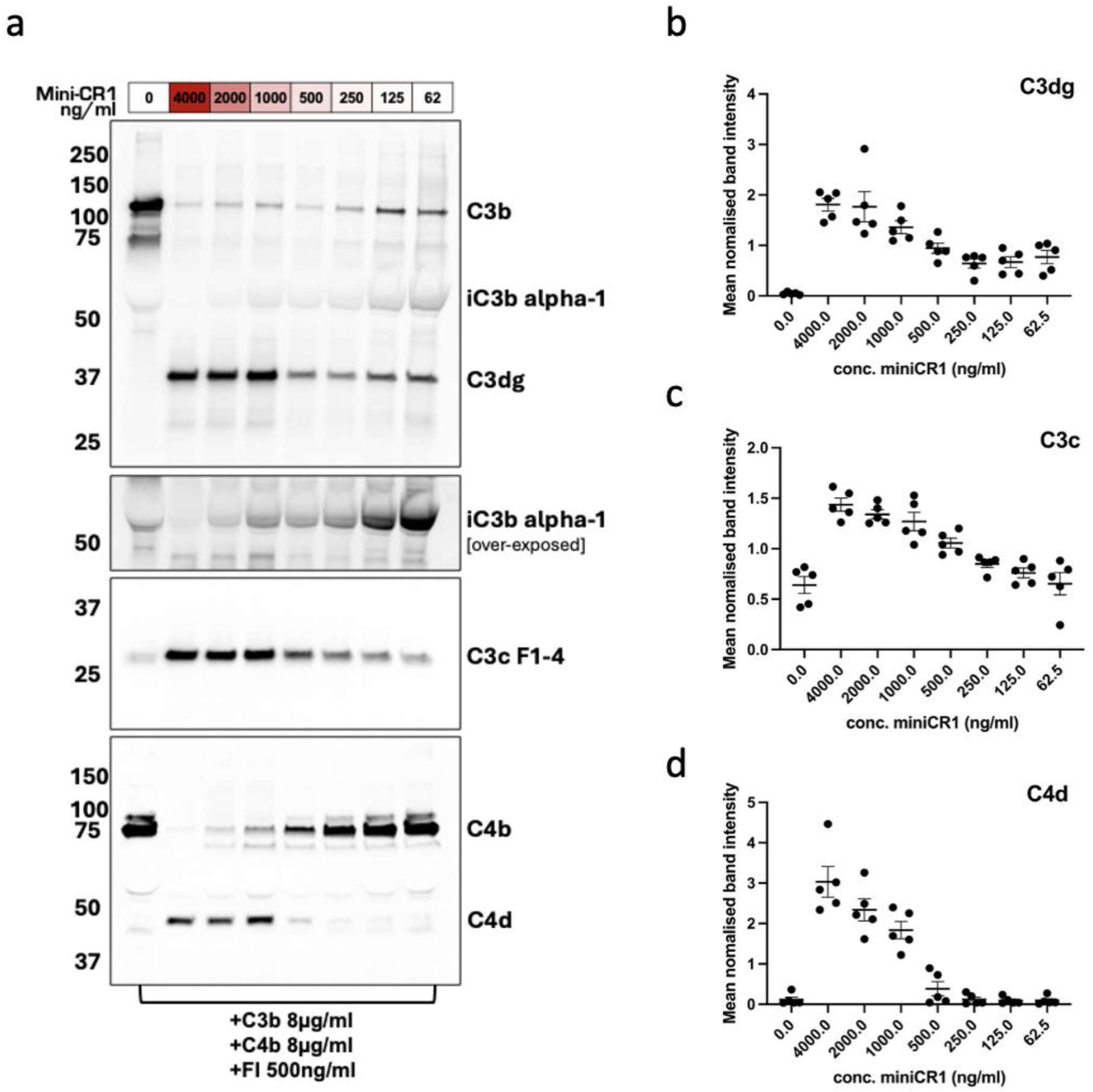
Recombinant mini-CR1 cleaves both C3b and C4b. a) Western blots assessing the bioactivity of mini-CR1 *in vitro* against high concentrations of human C3b 8 µg/ml and C4b 8 µg/ml, using a serial dilution of mini-CR1 ranging from 4000 ng/ml to 62 ng/ml (133nM to 2.1nM). A dose-dependent decrease in C3b, iC3b and C4b, with a corresponding does-dependent increase in their respective breakdown products, is shown: representative images of five separate experiments. Normalised densitometry values are plotted for the breakdown products C3dg (b), C3c (c) and C4d (d) showing the cumulative data from all five experiments ± SD.

Cleavage of C3b into iC3b and beyond by mini-CR1 would effectively inhibit the continuation of the amplification loop of the complement cascade. To confirm this, we used the alternative pathway Wieslab assay to assess the inhibitory effects on complement turnover in human serum; this was demonstrated with an IC_50_ of 125 nM with mini-CR1 (Figure 3A). Similarly, we tested serum from other species and saw for Cynomolgus monkey serum an IC_50_ of 26 nM, and rat serum 165 nM (Supplementary Figure 3). The binding affinity of mini-CR1 to immobilised C3b investigated by Biolayer Interferometry (BLI) was *Kd* = 2.1×10^-8^ M, stronger than for both FH (*Kd* 5.8×10^-7^ M) and FHL-1 (*Kd* 1.2×10^-6^ M); therefore, mini-CR1 has higher affinity for the C3b protein than other co-factors (Figure 3B). This replicates other studies that have found the C3b-binding regions of CR1 to have high C3b-binding affinities ^18,19^, and it is believed that this is the mechanism by which CR1 can drive the degradation of C3b by FI beyond iC3b ^20^.

**Figure 3:**
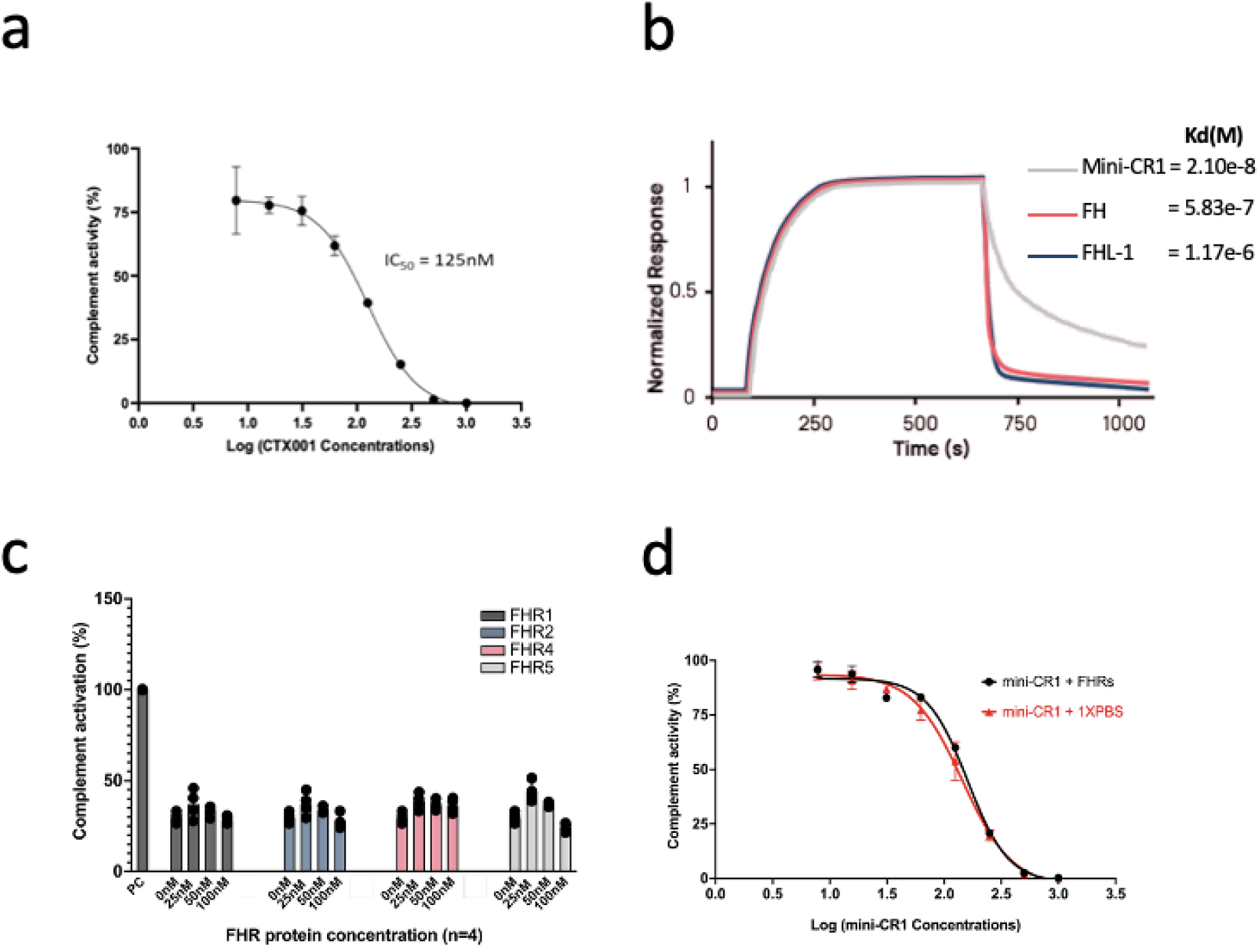
Further characterisation of mini-CR1 activity. a) Wieslab assay performed on zymosan activated human serum in the presence of increasing concentrations of mini-CR1. Mean (SD) are presented for three independent experiments. b) Biolayer Interferometry analysis of binding of fluid-phase mini-CR1, FH and FHL-1 to immobilised C3b demonstrating that mini-CR1 binds with the highest affinity. c) Complement activation levels in activated human serum (Wieslab assay) at fixed mini-CR1 concentration (161.7 nM) and increasing levels of supplemented FHR proteins. Mini-CR1 cofactor activity is not inhibited by any singular FHR protein tested, even at supraphysiological concentrations. Mean (SD) are presented, where: PC = Positive Control (human serum without CTx001 and without addition of FHR’s). d) Wieslab assay of complement alternative pathway activation in the presence of increasing levels of mini-CR1 and the presence (red) or absence (black) of physiological levels of FHR proteins: FHR1, 38 nM; FHR2, 55 nM; FHR4, 54 nM; and FHR5, 28 nM (29). IC50 in absence of FHR = 143.9 nM; IC50 in presence of FHR = 159.8nM, where data is shown as Mean (SD).

### Mini-CR1 is unaffected by increased levels of FHR proteins

FHR proteins have been demonstrated to be elevated in the circulation and/or retinal tissue in AMD and so provide a unique environment and an additional challenge for driving C3b breakdown by modifying co-factor activity. Supplementation of individual purified recombinant FHR proteins 1, 2, 4, and 5 (expressed and purified as described previously ^10^) into human serum to supra-physiological levels failed to interfere with the inhibitory effects of mini-CR1 on complement activation (Figure 3C). Similarly, when these FHR proteins were pooled and supplemented into human sera at levels matching the elevated concentrations found in AMD patients from previous studies ^9,11^ there was no effect on the IC_50_ value of mini-CR1, as shown in Figure 3D.

### Mini-CR1 can diffuse through human Bruch’s membrane

Complement turnover and MAC deposition associated with AMD occurs particularly on and around the ECM of the choriocapillaris layer underlying the retina ^5–8^. Bruch’s membrane alone, even without the retina, can present a barrier to therapeutics delivered to the retina (or vitreous) reaching the choriocapillaris. Therefore, we tested whether mini-CR1 can traverse Bruch’s membrane. Bruch’s membrane was enriched from five post-mortem human donor eyes using previously described protocols ^21^ and set within an Ussing chamber ^16^, which allows the passive diffusion of proteins from one chamber (sample chamber) into the other chamber (diffusate chamber) through the membrane being tested (Figure 4A). After 24 hours, for all five Bruch’s membrane samples, mini-CR1 is found in both chambers after passive diffusion through Bruch’s membrane, whereas other proteins tested including FH, FI and FB were not detected in the diffusate chamber after 24 hours (Figure 4B), as has previously been observed ^16^. C3b breakdown assays confirmed that mini-CR1 maintains its cofactor activity after crossing Bruch’s membrane, whereas pegcetacoplan (APL-2) struggles to diffuse through this tightly packed membrane^22^.

**Figure 4:**
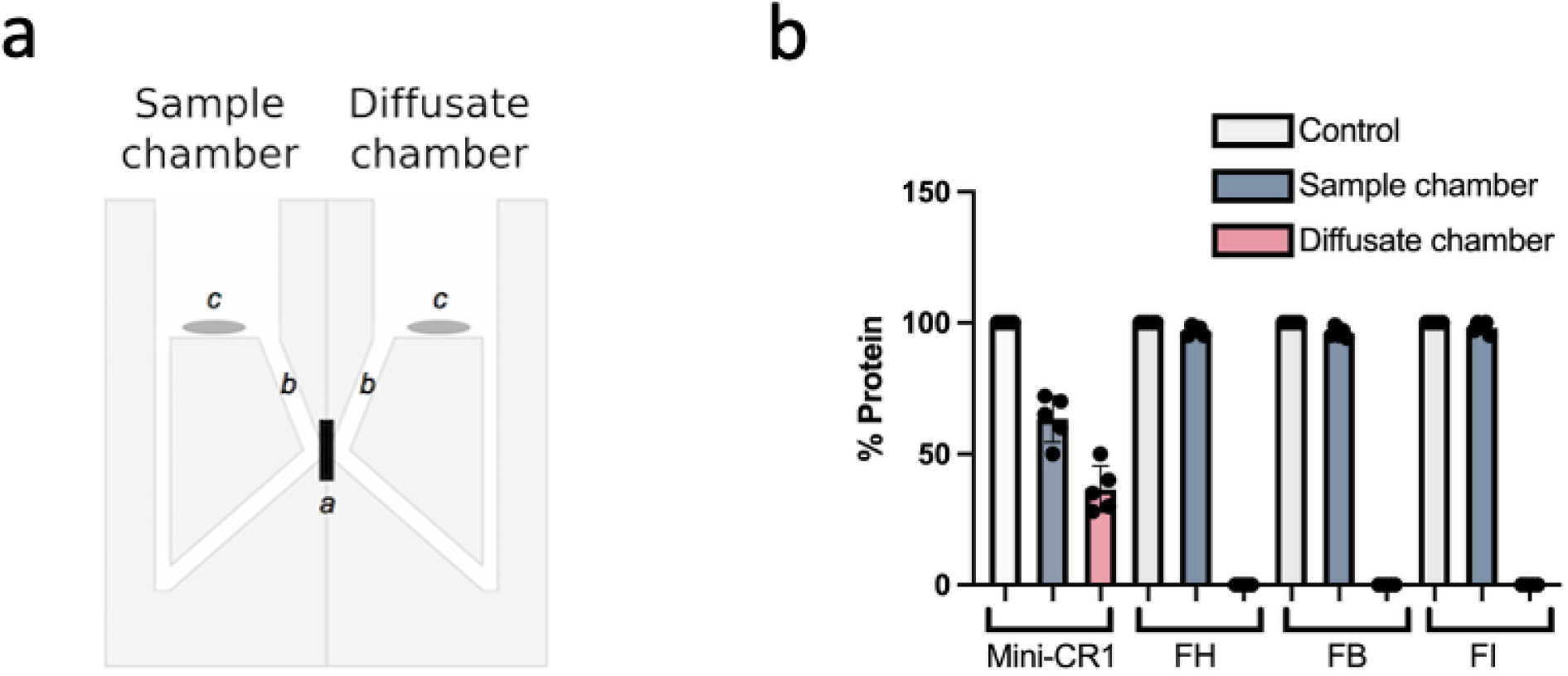
Bruch’s membrane diffusion properties of mini-CR1. a) Schematic representation of an Ussing chamber used in diffusion experiments, where a: enriched macula Bruch’s membrane; b: sampling access points; and c: magnetic stirrers. Recombinant mini-CR1, FH, FB or FI were placed in the sample chamber and after 24 hours the presence of protein in both chambers was assessed by SDS-PAGE. b) Summary analysis of diffusion experiments (Bruch’s membrane from n=5 AMD donors) analysed by SDS-PAGE showing that mini-CR1 is the only protein tested able to passively diffuse across Bruch’s membrane: mean (SD) depicted in the bar chart with individual data points shown.

### Vector design for rAAV delivery of mini-CR1 to RPE cells

Given the challenges associated with frequent intravitreal injections for patients with GA, we explored a gene therapy approach to give sustained delivery of mini-CR1. To achieve this, we designed and tested a series of different constructs for their ability to transduce the RPE and maximise mini-CR1 expression. We used *in vitro* and *in vivo* approaches (human RPE cell lines and naïve C57BL/6JRj mice following subretinal injection, data not shown) and on this basis took forward for further study a self-complementary design with rAAV2 serotype, already known to effectively target RPE cells ^23^, utilising a ubiquitous promoter and the secretory leader sequence from FH (Figure 5A). The self-complementary rAAV2 vectorised mini-CR1 (referred herein as CTx001) showed a dose-dependent increase in mini-CR1 transcription in RPE cells by qRT-PCR (Figure 5B). Subsequent transduction analysis using CTx001 at 100,000 MOI (multiplicity of Infection) on both RPE cell lines (hTERT-RPE1 and ARPE19) and human primary RPE showed mini-CR1 gene transcription by qRT-PCR (data not shown), and secretion of the mini-CR1 protein into the culture supernatant by Western blot analysis using a rabbit polyclonal antibody against mini-CR1 (Figure 5C).

**Figure 5:**
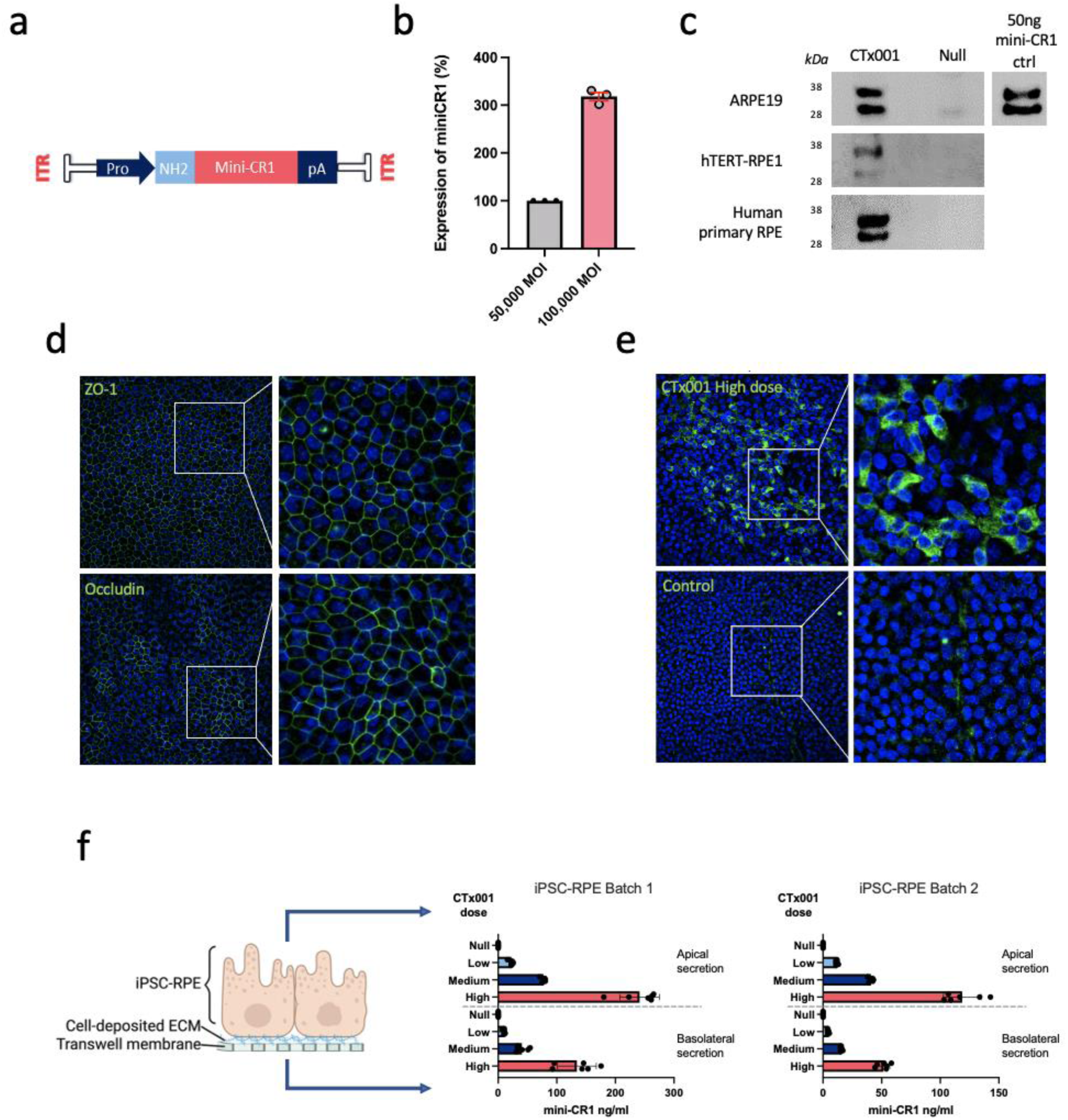
Transduction efficiency of CTx001 in RPE cell lines. a) Schematic of CTx001, a self-complementary rAAV2 expressing the mini-CR1 transgene with an N terminal secretory leader sequence. ITR = Inverted terminal repeat; Pro = Promoter; NH2 = Leader sequence; pA = Poly A sequence. b) Comparative analysis of transduction efficiency of the rAAV2 delivered CTx001 in driving mini-CR1 expression at 50,000 and 100,000 MOI in hTERT-RPE1 cells evaluated by qRT-PCR, 72 hours post-transduction. c) mini-CR1 expression and secretion from transduced human ARPE19 cells, hTERT-RPE1 cells and primary human RPE cells as assessed by Western blotting of culture supernatants. d) Quality control staining for iPSC-RPE tight junction formation showing both ZO-1 and occludin. e) Staining iPSC-RPE cells for mini-CR1 (green) after transduction with high dose of CTx001 (i.e. 2.2×10^10^ vg/transwell). f) Bi-directional secretion of mini-CR1 from iPSC-RPE cells seven days post transduction with CTx001 at one of three different doses: low, 9×10^8^ vg/transwell (∼8,000 MOI); medium, 4.5×10^9^ vg/transwell (∼40,000 MOI); or high, 2.2×10^10^ vg/transwell (∼200,000 MOI); compared to transduction with empty capsid (null) at 3.4×10^10^ vg/transwell as a control; mini-CR1 levels were measured using a Meso Scale Discovery (MSD) immunoassay. The results from two separate batches of iPSC-RPE are shown.

To investigate further the expression, and secretion, of mini-CR1 by RPE cells we employed induced pluripotent stem cell derived RPE cells (iPSC-RPE) that were fully differentiated in a transwell setting; these formed a confluent monolayer and expressed a series of RPE cell tight-junction markers, including occludin and ZO-1 (Figure 5D). These iPSC-RPE cells were transduced with three separate doses of CTx001 (low, 9×10^8^ vg/transwell; medium, 4.5×10^9^ vg/transwell; and high, 2.2×10^10^ vg/transwell); after seven days mini-CR1 staining was performed (Figure 5E) and samples taken from both the apical and basolateral chambers of the transwell system in which the cells were grown. Meso Scale Discovery (MSD) immunoassay showed secretion of mini-CR1 from transduced iPSC-RPE cells both apically and basolaterally (60% and 40% total secreted protein respectively), through the cell-deposited ECM and the transwell membrane itself (Figure 5F), thereby predicting effective delivery of mini-CR1 to both the neurosensory retina and through Bruch’s membrane to the choriocapillaris.

### CTx001 delivered mini-CR1 shows activity in RPE cell line

ARPE19 cells were transduced with CTx001 AAV2 (1.78 vg/cell) and maintained for seven days to allow stable expression, secretion and accumulation of mini-CR1 transgene protein. Cells were then incubated with 10% normal human serum to provide all necessary complement components for activation. Complement activation was triggered by addition of 100µg/mL zymosan and 10µg/mL LPS for 16h (to represent the long-term delivery of mini-CR1). LPS and zymosan promote complement activation by providing surfaces that facilitate C3b deposition and subsequent C5b9 formation: Zymosan activates the alternative pathway by serving as a binding surface for C3b, leading to amplification of the cascade^24^ and LPS engages both the classical and alternative pathways, further enhancing complement activation^25^. In both pathways, C3b and C4b contribute to the formation of C5 convertases (C3bBbC3b and C4b2aC3b), which cleave C5 into C5a and C5b. C5b then sequentially binds C6, C7, C8, and multiple C9 molecules, forming C5b9n (MAC), which mediates cell lysis. Mini-CR1, as a cofactor for FI, should augment C3b and C4b proteolysis, reducing C5 convertase formation and thereby limiting C5 cleavage and subsequent C5b9 assembly.

CTx001-AAV2 induced mini-CR1 expression in ARPE19 cells resulted in ∼700 ng/mL (∼24 nM) mini-CR1 in the supernatant as quantified by MSD (Figure 6A). Mini-CR1 secretion remained consistent regardless of the presence of human serum (grey bars) or serum supplemented with zymosan/LPS (red bars). The increase in mini-CR1 was accompanied by a significant reduction in C3b-iC3b1 levels in the supernatant (Figure 6B-C), measured using an MSD assay that first captures total C3b and then detects the 1H8 epitope, specific for C3b converted to iC3b^26^. These findings indicate that mini-CR1 in the ARPE19 supernatant actively facilitates the consumption of C3b and iC3b1 derived from human serum.

**Figure 6.**
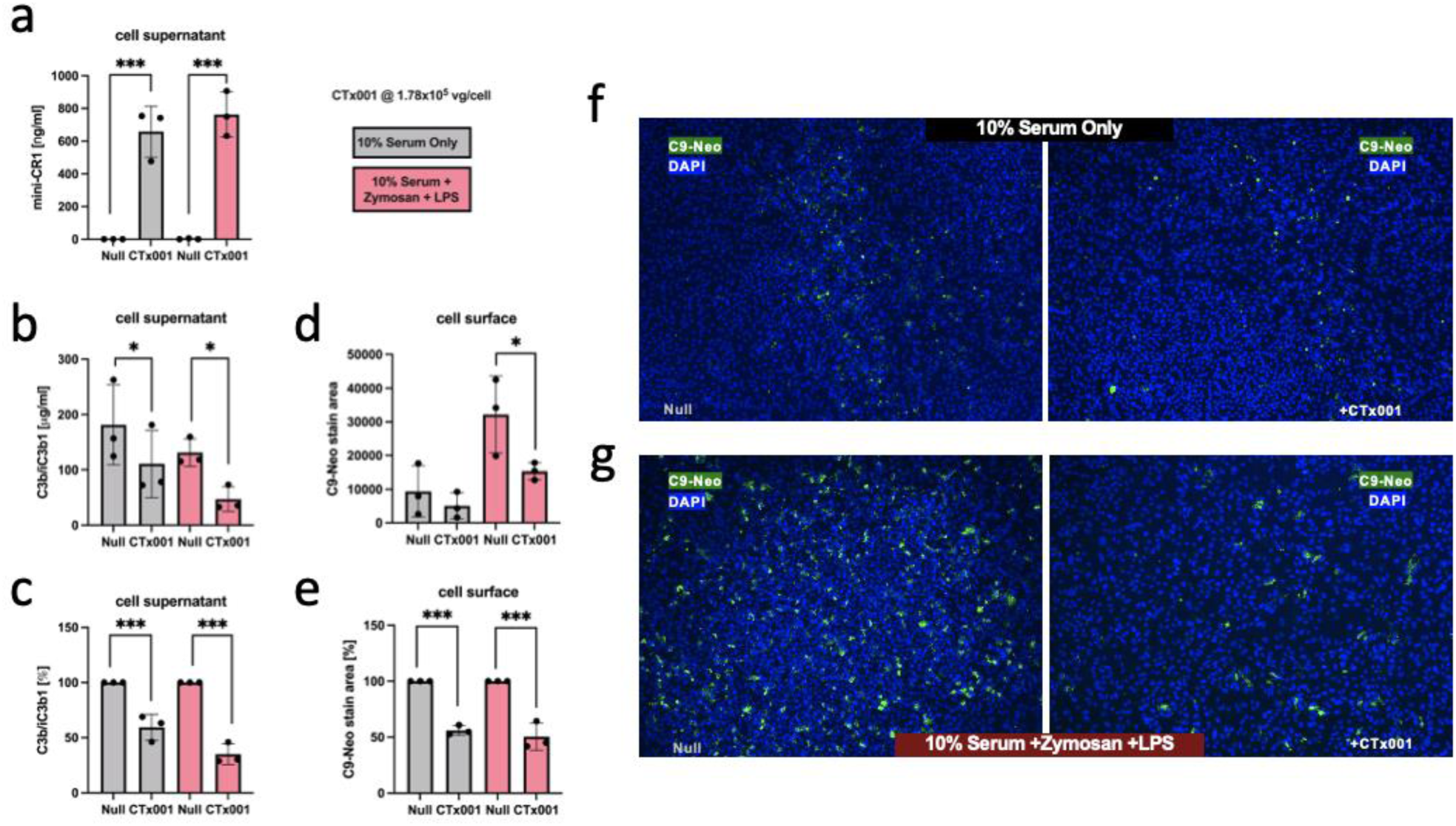
In vitro efficacy of mini-CR1 in RPE cell line transduced with CTx001. ARPE19 cells were transduced with CTx001 AAV2 at 1.78E5vg/cell] and maintained for 7 days in culture to allow for mini-CR1 transgene expression and protein accumulation in the culture medium. Control ARPE19 cells were incubated for 7 days with Null AAV2 (empty capsids) at 1.75E5vp/cell. a) soluble mini-CR1 was detected in the cell supernatants from ARPE19 cells incubated in either 10% normal human serum, or 10% normal human serum where complement had been activated by the presence of zymosan (100μg/m) and LPS (10μg/ml). b-c) The increase in mini-CR1 expression was accompanied by a significant decrease in C3b/iC3b levels in both normal and complement activated serum. d-e) C5b9n deposition on the surface of the ARPE19 cells was detected using a neoepitope-specific anti C9 antibody (clone aE11). A significant decrease in in C5b9n deposition on the cell surface is observed in the cells cultured with complement activated serum with cells transduced with CTx001. f-g) representative images of the C5b9n staining on cells cultured in either normal human serum (f) or complement activated serum (g). For each dataset, n=3 and bar graphs are plotted mean ± SD. Statistical comparisons were performed using ANOVA followed by Bonferroni’s post-hoc test to detect differences between control and CTx001-treated cells. * *p*<0.05 represents and *** *p*<0.005

Mini-CR1 bioactivity was further evaluated by staining for C5b9n on the surface of these serum- and zymosan-activated ARPE19 cells (Figure 6F-G). C5b9n deposition was detected using an aE11 neoepitope-specific antibody, which recognizes C9 incorporated into C5b9n complexes^27^. Incubation of ARPE19 cells with fresh human serum led to spontaneous accumulation of C9 neoepitope staining, reflecting MAC deposition on the cellular surface (Figure 6D and F). The addition of zymosan/LPS further increased C9 neoepitope staining, indicating enhanced C5b9n formation (Figure 6E and G). In CTx001-infected ARPE19 supernatants, where mini-CR1 reached ∼700ng/mL, a significant reduction was observed in C9 neoepitope staining, measured as cell surface deposits. Since C3b and all necessary complement components for C5b9n formation were provided by the addition of human serum, these results indicate that mini-CR1 not only reduces C3b/iC3b1 levels but also limits C5b9n (MAC) deposition on the cell surface.

### Subretinal delivery of CTx001 in a rat laser-induced CNV model

Having demonstrated that mini-CR1 inhibits complement by driving the complete degradation of C3b and the degradation of C4b, and that it can be delivered successfully to RPE cells with gene therapy, we aimed to test its efficacy *in vivo*. Rodent laser-induced CNV models were originally optimised for the study of ocular vascular diseases ^28^ and played a major role in the identification of VEGF as a target for ocular neovascularisation ^29^. Although not a model for GA in AMD (of which none exist), the laser CNV model results in increased complement turnover and the deposition of MAC within the retina ^30,31^. As such, complement inhibitors can be tested in this model ^32–34^, using MAC deposition as a biomarker of complement turnover in the posterior part of the eye.

It has been suggested that the mouse complement system does not behave in the same manner as that of human complement ^35,36^, and that in fact rat complement makes a far better model of the human system. Therefore, we performed *in vivo* efficacy experiments using a rat laser-induced CNV model. Subretinally delivered CTx001 at two different doses, low (1×10^8^ vg/eye) and high (5×10^9^ vg/eye), were compared to null vector (5×10^9^ vg/eye), with nine rats in each group (Figure 7A and Supplementary Table 1). Eye-cup lysates were collected 28 days after subretinal injections, and using the MSD immunoassay a dose-dependent secretion of the mini-CR1 protein was observed (Figure 7B). A reduction of 75.4% in the level of C5b9n staining was observed in rat eyes treated with high dose CTx001 compared to null vector alone (Figure 7C). Given that the raw data did not follow a normal distribution direct ANOVA analysis cannot be applied. However, log_10_ transformed data fulfil all of the tests for normality (Shapiro-Wilk, D’Agostino-Pearson, Kolmogrov-Smirnoff) and thus ANOVA analysis was applied and showed the 75.4% reduction to be statistically significant (*p*<0.01; Figure 7D).

**Figure 7.**
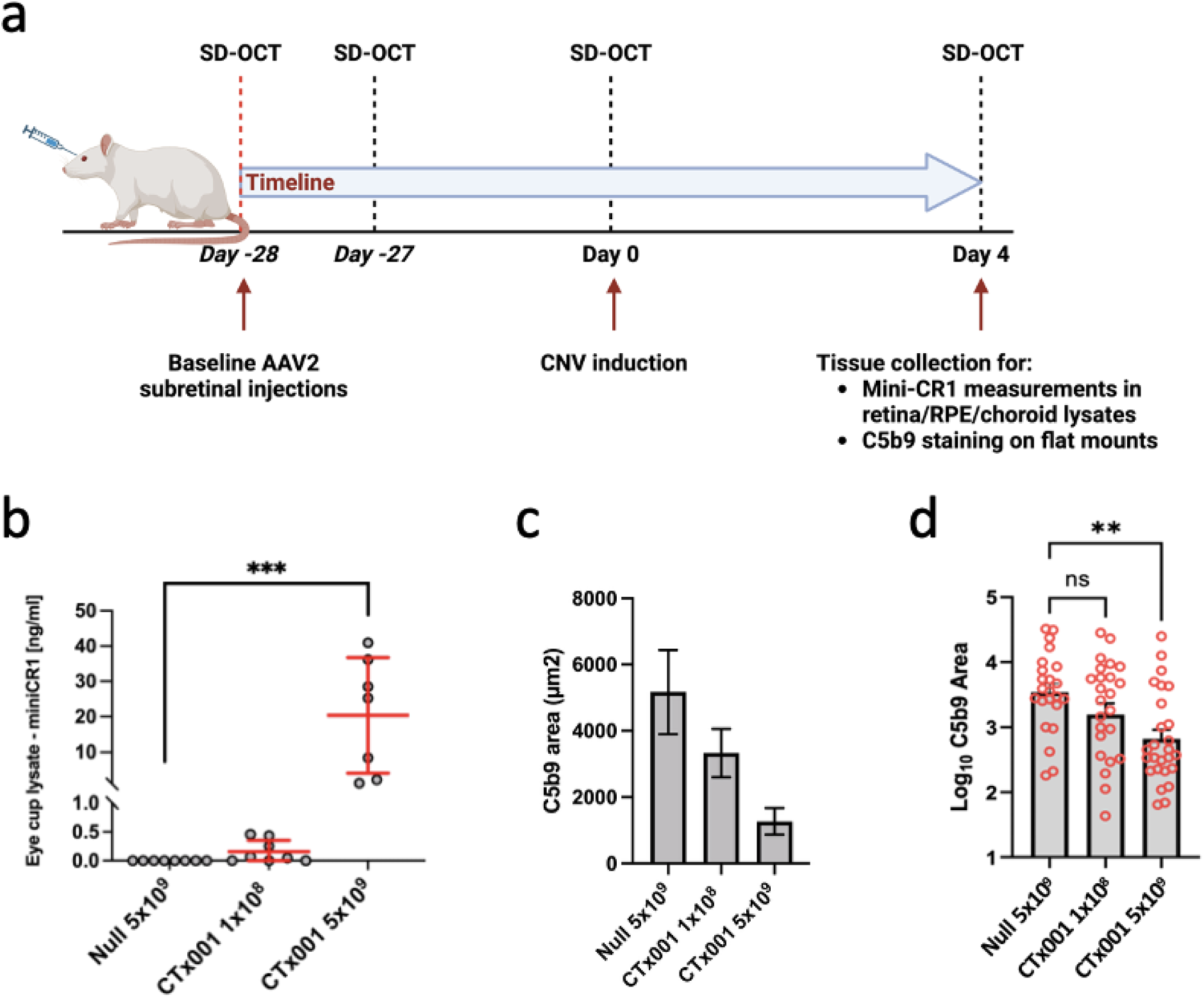
*In vivo* efficacy of CTx001-delivered mini-CR1 in a rat laser CNV study. a) Timeline schematic of the *in vivo* rat laser induced CNV model used in this study. The number of rats and details of each group are listed in Supplementary Table 1. Two different rAAV2 doses were injected: low, 1×10^8^vg/eye; and high, 5×10^9^vg/eye, n=9 rats per group. Control animals (n=9) received null rAAV2 subretinal injections (at 5×10^9^vg/eye). b) mini-CR1 protein expression was detected and quantified with a Meso Scale Discovery (MSD) immunoassay using eye-cup lysates. Representative images depicting laser lesion as defined by phalloidin staining and C5b9 staining within the lesion ROI after background threshold has been applied are shown in Supplemental Figure 4. c) C5b-9 deposition was measured by immunofluorescence in flat-mounted eyecups treated with null or CTx001, where mean values ± SD are shown. d) Log_10_ transformed data of the C5b-9 staining showing mean SD values; ANOVA with Dunnett’s post hoc was performed on log_10_ transformed data and demonstrated that high dose mini-CR1 group had a significant reduction in C5b-9 immunostaining (MAC formation). ** *p*<0.01 represents and *** *p*<0.005.

## Discussion

The first treatments approved for GA, pegcetacoplan and avacincaptad pegol, have validated the complement system as a therapeutic target. However, these require monthly or bimonthly intravitreal injections to maintain efficacy. This places considerable burden on both patients and healthcare providers and, while they slow GA lesion growth, they have not, yet, been shown to improve functional outcomes in visual acuity. Therefore, we elected to develop a complement modifier for treating AMD, CTx001, that is delivered as a single surgical treatment and produces a sustained effect, and which targets multiple pathways involved in complement activation and turnover. CTx001 is a gene therapy designed to be delivered by subretinal injection (i.e. creation of a transient bleb between the neurosensory retina and RPE). As it uses an rAAV2 vector, it is expected to predominantly target the RPE ^37^, which then secretes the small, soluble complement inhibitor protein, mini-CR1, in a so-called bio-factory approach.

We demonstrated that RPE cells transduced with CTx001 secrete mini-CR1 apically and basolaterally (Figure 5F). Furthermore, we demonstrate that it inhibits MAC formation on the surface of CTx001 transduced ARPE19 cells and promotes C3b consumption in the cell culture media (Figure 6). Using human Bruch’s membrane *ex-vivo*, we demonstrated that mini-CR1 crosses Bruch’s membrane (Figure 4); it is important to test this using human tissue as there is wide variation in Bruch’s membrane thickness and permeability between species ^38^. The ability of an AMD therapeutic to cross Bruch’s membrane is now recognised as an important consideration for therapeutic design and delivery modality selection ^39,40^. Bruch’s membrane acts as a selective barrier to many proteins, including complement proteins, based on size, hydrodynamic radius, and charge ^16,41^; if complement therapies are delivered to the vitreous or retina, failure to cross Bruch’s membrane may pose a significant barrier to realising their full efficacy potential, as their function will be limited to retinal tissues and they will not treat complement over-activation in the choriocapillaris layer. Therefore, we predict that the transduced RPE cells will continually secrete mini-CR1 that will reach both the retina and choroid, thereby inhibiting complement over-activation in all anatomical compartments where complement over-activation has been demonstrated in AMD.

We have shown that mini-CR1 is a potent cofactor for the FI-mediated degradation of C3b, iC3b and C4b, so it targets the alternative and classical complement pathways. The breakdown of C3b to iC3b removes it from the complement C3 amplification loop, thereby decreasing anaphylatoxin production and downstream MAC deposition. Mini-CR1 is an effective co-factor for FI and is more potent than other co-factors including FH and FHL-1, so it can breakdown C3b beyond iC3b to C3dg. This can be explained by its higher affinity to C3b compared to FH and FHL-1 (Figure 3B). Whilst iC3b cannot contribute to the C3 amplification loop, it does act as a strong ligand for CR2, CR3 and CR4 receptors on immune cells, and the resultant interaction with immune cells results in continued opsonisation and remains pro-inflammatory. The further degradation to C3dg by mini-CR1 removes, or greatly reduces, interactions with immune cells, thereby further reducing their potency and the pro-inflammatory microenvironment.

Recently, genetically driven elevated levels of circulating FHR proteins have been associated with AMD pathogenesis ^9–11^. Indeed, there is evidence from Mendelian randomisation studies that high FHR protein levels cause AMD. These highly homologous proteins are closely related to the FH protein and share some of its binding partners ^42^. FHR proteins bind to C3b but cannot bind to FI, so they are not cofactors for FI-mediated C3b breakdown; instead, they can act as complement activators by outcompeting FH and/or FHL-1 binding to C3b, thereby decreasing C3b breakdown and causing excessive complement activation ^10,43^. Given that these FHR proteins accumulate in the ECM surrounding the choriocapillaris in human eyes, as well as in drusen within Bruch’s membrane ^10,11,44^, we assessed whether FHR proteins hinder the co-factor activity of mini-CR1. We demonstrated that mini-CR1 is not displaced by supra-physiological levels of FHR proteins in human serum, so mini-CR1 maintains cofactor activity for FI in the presence of FHR proteins (Figure 3D-E).

There are no satisfactory *in-vivo* models of GA. However, in rodent laser choroidal neovascularisation (CNV) models complement overactivation is observed. This occurs in the region of the retina and choroid, so it is in an anatomically relevant location. We used the rat laser CNV model, and the readout used for the effect of CTx001 was deposition of the terminal complement activation product MAC. We demonstrated significantly reduced levels of MAC deposition following laser CNV induction (75% compared to null vector controls; Figure 7). MAC deposition has also been used to evaluate soluble CD59, the native complement regulator that naturally targets MAC cellular deposition directly ^45^, as a potentially therapeutic complement modifier in the eye. In this case, MAC deposition in the laser-induced CNV mouse model is reduced by ∼50% ^32^. Other groups have used CNV lesion size and leakage as a readout, but we believe MAC is a more direct and specific readout of complement overactivation.

In summary, we present here a first generation, self-complementary rAAV2 mediated gene therapy (CTx001) that successfully transduces RPE cells resulting in secretion of the soluble complement modifier mini-CR1. This modifier can address both complement-meditated amplification and opsonisation, in retinal tissues, Bruch’s membrane and the choriocapillaris. This ‘one shot and done’ delivery approach, was shown to be very efficacious in reducing complement turnover and can target the complement cascade at the levels of both C3b, iC3b and C4b. CTx001 aims to address the continued unmet need in the therapeutic landscape for intervention in GA.

Supplementary information accompanies the manuscript on the Experimental & Molecular Medicinè website (http://www.nature.com/emm/)

## Data Availability Statement

The data that support the findings of this study are available from the corresponding author(s) upon reasonable request.

## Acknowledgements

Medical writing and editorial assistance in the development of this manuscript was provided by Jennifer Green, PhD (Green Ink Communications Ltd, UK), funded by Complement Therapeutics. SJC receives financial support from the Helmut Ecker Foundation. We thank the staff at The University of Manchester Biomolecular Analysis Core Facility for support with the BLI experiments, and Powered Research, USA who performed the rat studies. This work was partially funded by the Medical Research Council, UK (MR/K024418/1), and was facilitated by the Manchester NIHR Biomedical Research Centre and the Greater Manchester Local Clinical Research Network.

## Author contributions

Project conception and initiation: SJC, PNB, RDU, MM, DK, RH; designing research studies: SJC, PNB, RDU, MM, DK, RH, SR; conducting experiments: SR, AD, SP, RG, PEB, NP, PW, VT, ES; acquiring data: SR, AD, SP, RG, PEB, NP, PW, VT, ES; analysing data: SJC, PNB, RDU, MM, DK, RH, SR, AD, SP, RG, PEB, NP, PW, PZ, VT, ES; providing reagents: SR, VT, ES; writing the manuscript: SJC, PNB, RDU, MM, DK, RH, SR, AD, SP, RG, PEB, NP, PW, PZ, VT, ES.

## Declaration of Interests

SJC, PNB, RDU are co-founders of Complement Therapeutics (CTx), hold founding shares, and are named on patents associated with this work. MM, AD, SP, RG, PEB, NP, PW, PZ, RH, DK are employees of CTx. MM, RH, DK, PEB hold equity in CTx. SR, VT and ES have no conflicts of interest.

## Supplementary Information

### Supplementary Methods

#### Purified recombinant mini-CR1 protein production in HEK293T cells

1 mg/ml plasmid DNA and 7.5 mM PEI max transfection reagent (Polysciences, Germany) were added to separate aliquots of 150 mM NaCl and each incubated for 10 min at room temperature. The DNA/PEI mixture was added to a 15 cm diameter dishes containing 7×10^6^ HEK293T cells/per dish in a dropwise manner and incubated at 37°C. After 5–6 hours, the media was replaced with 2% (v/v) FCS containing DMEM (no antibiotic) and incubated at 37°C for overnight. Conditioned media was collected daily and replaced with fresh media. Conditioned media from each of 20 dishes (17.5 ml/dish) were collected and pooled after 24, 48, 72, and 144 hours. The pooled conditioned media after 144 hours (1,400 ml total) was diluted with the addition of 600 ml 50 mM HEPES, 500 mM NaCl, 20 mM Imidazole, pH 7.5. To this, 12 ml of NiNTA resin (Expedeon) was added and incubated overnight with rotation at 4°C. The NiNTA beads were collected by passing the media through empty PD10 columns with a filter (GE Healthcare) by gravity flow (500 ml media per PD10 column, i.e. four PD10 columns in total). The beads were then washed with 10 ml of wash buffer (50 mM HEPES, 500 mM NaCl, 20 mM imidazole, pH 7.5). Finally, the His-tagged mini-CR1 protein was eluted using 16 ml of elution buffer (50 mM HEPES, 500 mM NaCl, 500 mM imidazole, pH 7.5). Eluted protein was dialysed back into wash buffer overnight at 4°C before being concentrated further by the addition of 1.5 ml NiNTA beads and then eluted into 6 × 1 ml aliquots. Purified protein aliquots were dialysed into 20 mM glycine, 125 mM NaCl, pH 9.0 using a Slide-A-Lyzer dialysis cassette (Thermo Fisher Scientific, USA) with a 10 kDa cut off. The purified recombinant protein was assessed for purity by SDS-PAGE and visualised by staining the gels for 60 min at room temperature with Instant Blue Coomassie stain (Expedeon, UK).

#### C3b/C4b degradation assays

The assay mixture contained C3b (8 µg/mL), C4b (8 µg/mL), and FI (0.5 µg/mL), all from CompTech, USA, diluted in 1× PBS with 500 µg/mL BSA. Mini-CR1 was serially diluted from 4000 ng/mL (133 nM) to 62 ng/mL (2.1 nM). Reactions were incubated at 37 °C with shaking (800 rpm, Thermomix) for 16 hours (x2 predicted half-life of mini-CR1) to model the sustained exposure expected with *in vivo* gene therapy approach. Following incubation, samples were prepared for immunoblotting. A total of 120 µL from each reaction was mixed with 40 µL of 4× Laemmli SDS sample buffer, boiled at 95 °C for 10 minutes, and 25 µL was loaded per lane. Proteins were separated on Bolt™ Bis-Tris 4–12% protein gels and transferred nitrocellulose membranes. Membranes were blocked in 10% fat-free milk powder dissolved in 1× PBS with 0.05% Tween-20 (PBS-T) for at least 60 minutes, then incubated overnight at 4 °C with primary antibodies. The following primary antibodies were used: rabbit anti-C3d (Dako), rabbit anti-C4d (Abcam), and mouse anti-C3c clone F1-4 (Hycult), each diluted 1:500 in 5% BSA in 1× PBS. After primary incubation, membranes were washed four times for 10 minutes each with PBS-T. HRP-conjugated swine anti-rabbit IgG (1:2000 in 5% BSA) was applied for 65 minutes, followed by four additional washes. HRP signals were detected using ECL-Plus reagents (2-minute incubation) and visualized on the iBright 1500 imaging system. Immunoblots were quantified by densitometry of chemiluminescence signals using ImageJ for C3b, C3dg, C3c, C4b and C4d. For each of the five independent C3b/C4b digestions and corresponding immunoblots, raw density values were scaled by mean normalization. Specifically, each lane’s value was normalized to the average densitometry value of all lanes within the same blot.

#### Alternative complement pathway inhibition assays

Mini-CR1 protein at concentrations ranging from 1000 nM to 7.8135 nM was prepared via 8-fold serial dilution in 200 μL of the assay buffer. For the positive control (PC) Zymosan activated human serum and cynomolgus monkey serum (diluted 1:18) and rat serum (diluted 1:5) without the addition of mini-CR1 were used, while dilution buffer alone or serum without complement activity were used as the negative control (NC). Subsequently, 100 μl of each (1000 nM to 7.8135 nM) mini-CR1 protein was added to 100 μl of 1:18 pre-diluted activated human serum and cynomolgus monkey serum and 1:5 pre-diluted rat serum in the assay diluent and allowed to incubate for 30 minutes at RT. After incubation samples, PC, NC, and diluent alone (Blank) were transferred to the assay plate in duplicate and further incubated for 1 hour at 37°C followed by washing three times with the 1x wash buffer. Next, 100 µL of the conjugate/tracer, was added to each well and incubated for 30 min at RT. The conjugate in the Wieslab assay consists of an alkaline phosphatase-conjugated antibody targeting the primate C5b-9 neoantigen, while the Hycult assay uses a biotinylated antibody specific to rat C5b-9. Therefore, for detection of the bound antibodies in the Weislab assay, an alkaline phosphatase substrate solution (PnPP) was used, 100 μL in each well, and incubated for 30 min at RT. While in the Hycult assay, 100 μL of streptavidin peroxidase solution was added, incubated for 30 min at RT followed by washing and 100 μL of TMB in each well. After 30 mins incubation in the dark, the reaction was stopped by the addition of oxalic acid solution. The amount of complement activation was measured by the absorbance at 405 nm (Weislab assay) or 450 nm (Hycult assay) using a Molecular Devices VersaMax Microplate Reader (Marshal Scientific, USA). The results were expressed as a percentage of MAC (C5b-9) formation and calculated using the formula: (Sample-NC)/(PC-NC) x100. IC_50_ values were analysed in GraphPad Prism with inhibitor versus response analysis employing a four-parameters variable slope fit.

### Binding affinity studies using Biolayer Interferometry

Measurements of binding affinities for mini-CR1, FHL-1, and FH to C3b were performed using an Octet RH384 Biolayer Interferometry (BLI) system (Satorius, Germany) with biotinylated C3b immobilized onto streptavidin biosensors. Experiments were performed at 25°C and in PBS with 0.2% (v/v) surfactant P20 (PBST). Biotinylated C3b was coated onto Streptavidin biosensors in a total volume of 200 μl/well, at a concentration of 1 mg/ml for 10 minutes. Each protein of interest (i.e. mini-CR1, FHL-1 or FH) was tested separately, in triplicate, in concentrations ranging from 50-1000 nM with the following data acquisition settings: loading, 600 sec; baseline, 150 sec; association, 600 sec; and dissociation 600 sec. After subtraction of control (blank) sensor values for each condition, association and dissociation rate constants were determined by global data analysis using the Octet® data analysis software.

#### Ussing Chamber Diffusion

In each experiment a 5 mm diameter disc of macular Bruch’s membrane obtained from human donor eyes without macroscopic evidence of AMD or other macular pathology was mounted in an Ussing chamber, forming a barrier between two identical compartments as shown in Figure 4A. Both sides of the Bruch’s membrane were subjected with a 5 min wash of 2 mL PBS at room temperature. PBS (2 mL) or purified proteins recombinant mini-CR1, FH, FB or FI in PBS (100 µg/mL, 2 mL volume) were added to the sample chamber. After 1 min, if no leaks occurred into the second compartment, 2 mL PBS was added to the diffusate chamber. This set up was maintained at room temperature for 24h with gentle stirring in both compartments to prevent protein diffusion gradients. Subsequently, 20 µl samples from each chamber were analyzed by gel electrophoresis. Gels were either stained with Instant Blue stain or subjected to Western blotting. To calculate the percentage of protein in the sample or diffusate chambers, band densities in the Instant Blue stained SDS gels were measured using ImageJ64 (version 1.40g; http://rsb.info.nih.gov/ij). The average intensity of these bands, over five separate experiments, were compared to the density of control bands that represent 100% loaded protein (i.e., 20 µl of 100 µg/ml). The calculated percentage protein was then plotted ± SD. For checking cofactor activity of mini-CR1 after diffusion, C3b breakdown assay was performed using 2 µg of C3b and 0.04 µg FI with either FH or 10 µL sample taken from diffusate chamber as described above.

#### Transduction of RPE cell lines and analysis of expression

For the transduction of ARPE19, hTERT-RPE1, and primary RPE cells, 90,000 each of these cells were plated in 24-well plates and after 24 hrs cells reached 50% confluency i.e. 120,000 cells/well. Once the cells reached 50-60 % confluence, the culture medium from cells was removed and 500 μL of medium containing either CTx001 or null vector rAAV were added, at multiplicity of infections (MOIs) ranging 10,000, 50,000 and 100,000. These viral transduction rates experiments were performed in triplicate. After 72hrs post-transduction, media was collected, PMSF was then added (1.5 mL media + 1.2 μL PMSF) to the conditioned media which was subsequently analyzed by Western blotting. Additionally, cells were collected for qRT-PCR analysis to further evaluate the transduction efficiency.

Western blotting was performed to validate the transduction efficiency of rAAV2 expressing mini-CR1 (CTx001) in RPE cell lines and for evaluating cofactor activity of mini-CR1 after diffusion through Bruch’s membrane (from Ussing chamber experiment). For Western blotting, a wet transfer apparatus was used applying 20V for 1 hour, and the protein were transferred onto PVDF membrane in a buffer containing 25 mM Tris, 192 mM glycine, 10% methanol. PVDF membranes were blocked in 10% milk, 0.2% BSA in PBS overnight at 4°C before addition of polyclonal rabbit anti-human mini-CR1 antibody (1:500 diluted, produced in-house) against mini-CR1 protein and anti C3 antibody (clone 1H8, Abcam, UK) for detecting C3-breakdown products for 1 hour at RT. Membranes were washed three times for 30 min each in PBS-T (0.2% tween 20 in 1x PBS) before the addition of HRP conjugated secondary antibody for 1 hour at RT. Membranes were washed three times for 30 min each in PBS-T (0.2% tween 20 in 1x PBS) before the addition of 40:1 ECL Plus Western blotting substrate (Pierce, Thermo Fisher) for 1 min in the dark and then the Fusion imager was used for detection of protein bands.

For rtPCR RNA was extracted from rAAV-mini-CR1 construct transduced and untransduced ARPE19 cells, hTERT-RPE cells, and primary RPE cells using the Isolate RNA Mini Kit (Bioline, catalogue number BIO-52072), and from the posterior cups of rats using Total RNA Purification Plus Kit (Norgen Biotek) following the respective manufacturer’s protocol. Isolated RNA was quantified using nanodrop and converted into cDNA using the Superscript IV Vilo Master Mix (cat no. 11756050, Invitrogen). Quantitative PCR was performed using custom-designed specific mini-CR1 Pair 1 FAM-labelled TaqMan probes, (Forward Primer Sequence: AGGACGTGTGCAAGAGAAAG, reverse primer sequence: GTGGTGCAGCTGTAGTTGAT, Probe sequence: CTCCTGATCCTGTGAACGGCATGG). In brief, 10-100 ng of cDNA was suspended in a reaction mix consisting of 1 µl of Mini-CR1 Pair1 TaqMan probe (target) and either 1 µl of VIC labelled TaqMan GAPDH probe (Hs02758991_g1) or 18s probe as control, 10 µl of 2x reaction TaqMan™ Universal Mastermix II, no UNG (Applied Biosystems™, ThermoFisher Scientific, cat no. 4440040), in a final reaction volume of 20 µl in 96 well plates. The qRT-PCR samples were run in triplicate in an ABI 7500 Real-Time PCR system (Applied Biosystems) under the following thermal cycling conditions: initial denaturation at 95°C for 10 min, followed by 40 cycles of denaturation at 95°C for 15 sec and annealing/extension at 60°C for 1 min. Gene expression levels were normalized to *GAPDH* or 18s expression and the relative expression was determined by the ΔΔCt method.

#### Human iPSC-RPE culture and transduction

Human induced pluripotent stem cell (iPSC) derived RPE cells (iPSC-RPE, Phenocell, France) were cultured according to manufacturer’s instructions. Briefly, cells were thawed and then seeded onto Matrigel® coated (final density of Matrigel 8-10 µg/cm^2^) transwell-12 inserts at a density of 100,000 cells/cm^2^. The cells were then grown for 28 days in culture medium containing 70% DMEM, high glucose, 30% Ham’s F12 Nutrient Mix, 2% B-27® Supplement, 1% Antibiotic-Antimycotic; by which time the cells acquired their characteristic polygonal morphology and were pigmented. This maturation was confirmed also with ICC staining for occludin and ZO-1 proteins. Transwells containing iPSC-RPE (each ∼112,000 cells) were transduced with one of three doses: low, at 9×10^8^ vg/transwell (∼8,000 MOI); medium, at 4.5×10^9^ vg/transwell (∼40,000 MOI); or high, at 2.2×10^10^ vg/transwell (∼200,000 MOI). An empty capsid, null control was also included at 3.4×10^10^ vp/transwell. The transduction solutions, 0.8 ml apical side (insert) and 0.8 ml basolateral side (bottom well), total 1.6 ml per cell system, were incubated for 3 days. After 3 days of transduction, apical (∼0.7 ml) and basolateral (∼0.7 ml) media samples were collected in tubes containing protease inhibitor cocktail on 3, 7, 10, 14, 17 and 21 days. After the media sample collection, fresh cell culture media were added in the apical and basolateral side of the cells and samples were frozen and stored at -70°C. These experiments were repeated to test consistency between batches of cells.

#### Immunostaining of occludin, ZO-1 and mini-CR1

Methanol fixed cells were washed with 1x DPBS and blocked with 10% (v/v) normal goat serum in DPBS for 20 min. Rabbit anti-occludin, rabbit anti-ZO-1 and a custom anti-human mini-CR1 rabbit monoclonal antibody (FairJourney Biologics, Porto, Portugal, clone FJ2309) primary antibodies were diluted (1:100) in 0.5% normal goat serum in 1x DPBS and incubated at 2-8°C overnight. Cells were washed carefully with 0.5% normal goat serum in 1x DPBS and immunoreacted with anti-rabbit secondary antibody AF488 (1:500) for 90 min. Cells were washed carefully with 0.5% goat serum in 1x DPBS and nuclei were stained with 4′,6-Diamidino-2-phenylindole dihydrochloride (DAPI, 1:5000) for 5 min. Cells were cut-out with the membrane and placed on microscope slide, mounted with Fluoroshield and cover slipped. The no primary antibody control was included in the experiments. Signals from ZO-1, occludin and mini-CR1 were captured at 40X in the green channel and DAPI in the Blue channel using a Leica K5 camera (Leica Microsystems GmbH) attached to a Leica Thunder Imager 3D Tissue (DM6B-Z, Leica Microsystems GmbH). The whole system was controlled by the LAS X software (v. 3.7.5.24914, Leica Microsystems GmbH).

#### Mesoscale [MSD] electrochemiluminescence Mini-CR1 and C3b-iC3b quantitation assays

MSD assays were performed to assess Mini-CR1 and C3b-iC3b in ARPE-19 cell lysate or culture medium supernatant samples. Standard 96-well MSD plates were coated overnight at 4°C with the appropriate capture antibodies diluted to 8 µg/mL in 1× PBS: a custom anti-human mini-CR1 rabbit monoclonal antibody (clone FJ225_mAb02F09, FairJourney Biologics) for Mini-CR1 assessment, and a goat polyclonal anti-human C3 antibody for C3b-iC3b measurement (Invitrogen). Following incubation, plates were blocked with 5% BSA in 1×PBS on a plate shaker (750 rpm) for 90 minutes at room temperature, then washed three times with 1× PBS. Standard curves were prepared using purified human Mini-CR1 (125-0 ng/mL), and C3b (500–0 ng/mL), all diluted in 500 µg/mL BSA in 1×PBS. Blank wells contained 500µg/mL BSA in 1× PBS. Samples and standards were added to the plate, sealed, and incubated on a plate shaker (750rpm) for 90 minutes at room temperature, followed by three washes with 250µL 1× PBS. For detection, monoclonal antibodies were sulfo-tagged using manufacturers’ protocol (MSD, Rockville, US): a custom anti-human mini-CR1 rabbit monoclonal antibody (clone FJ225_mAb03H10, 2 µg/mL, FairJourney Biologics), and an anti-human C3b-iC3b1 antibody (clone 1H8, 1 µg/mL, Abcam, UK), all diluted in 500µg/mL BSA in 1× PBS. Plates were incubated with detection antibodies for 60 minutes at room temperature, followed by four washes with 1× PBS-Tween 20 (0.05%), and a final wash with 1× PBS. Just before analysis, MSD reading buffer was prepared by diluting MSD Reading Buffer 1:1 in DI water then added to the plate and immediately analyzed using the Meso QuickPlex SQ reader. Data were processed using MSD Discovery Workbench software, applying standard analysis parameters.

#### C9 Neoepitope staining

Supernatants were carefully removed and wells were washed with PBS. After discarding PBS, absolute ethanol was added to each well and incubated for 10min to fix the cells. The wells were subsequently washed twice with PBS. Following washing blocking buffer—composed of 10% normal donkey serum in 5% bovine serum albumin—was added to each well and incubated for 1h at room temperature to prevent non-specific antibody binding [secondary antibodies are raised in donkey]. The blocking buffer was then discarded, and the wells were incubated overnight at 4°C with primary antibody solution [mouse anti-human TCC C9 neoepitope antibody; clone aE11; HyCult]. Following overnight incubation, wells were washed three times with PBS. Next, secondary antibody [donkey anti-mouse Alexa-fluor 488] was added to each well and incubated for 75min at room temperature. Next, secondary antibody solution was removed, and wells were washed three times with PBS for 10min. Finally, the wells were covered with Fluoroshield Mounting Medium [containing DAPI to counterstain nuclei].

#### C9 Neoepitope staining imaging and quantitation

For consistency, multiple 10x images per well were captured [eVOS instrument], with images taken from the top, middle, left, and right areas of each well. Immunofluorescence signal quantitation was performed using ImageJ. Each imaging channel from the 10x images was transformed into 8-bit format. Background staining was subtracted individually for each image, ensuring consistent baseline correction across all fluorescence channels. Staining quantitation was performed using ImageJ’s thresholding function to enhance consistency. Image acquisition settings varied depending on the channel: for DAPI, the light intensity was lower than 10 and exposure ranging between 9 and 90ms, while for 488 [C9 neoepitope], the light intensity was set to 100, with exposure at 350ms. Digital gain was not used to reduce quality artefacts. Upon image input, channels were split, and the 488 channel, corresponding to C9 neoepitope staining, was used for analysis. The image was processed in 8-bit format, and background removal was performed by disabling smoothing with a rolling ball radius of 50 pixels. After background subtraction image threshold was adjusted to 20 for all images to provide for consistent and comparable quantitation across images, conditions and plates.

#### Staining of choroidal flat mounts

Choroidal flat-mounts were incubated overnight with fluorescein-labeled isolectin B4 (from Griffonia simplicifolia lectin I) (dilution 1:200) to detect neovascularization and subsequently incubated overnight with rabbit C5b-9 antibodies (dilution 1:350) to detect MAC (membrane attack complex) deposition. Following this, choroids were incubated for an additional 3 hours with goat anti-rabbit antibodies conjugated with AlexaFluor 594 fluorescent marker to visualize primary C5b-9 antibody. After each incubation step, samples were thoroughly washed for 10 mins with TBS (Tris-buffered saline) buffer to remove any unbound antibodies and other contaminants. Samples were then mounted using Fluoroshield. Slides were imaged using a Leica B6 microscope (Leica Microsystems). The stained areas were outlined and measured using a validated protocol in the image processing software FIJI. Each image was evaluated individually, by identifying lesion and thresholding.

---

**Supplementary Table 1:** Experimental groups used in the *in vivo* rat laser-induced CNV study.

| Group | Treatment | # Animals<br>/ #<br>injected<br>eyes (OD<br>injected) | CNV(OD) | Dose (vg/eye) | Dosing<br>Volume | FA and<br>OCT | Flat mount<br>(OD eyes) |
| --- | --- | --- | --- | --- | --- | --- | --- |
| 1 | Null | 9 rats/18<br>eyes | Yes | 5x10<sup>9</sup>vg/eye,<br>subretinal | 2.5 µL | At day 0<br>and day 1 | N=10<br>eyes/grp for<br>flat mounts<br>N=8<br>eyes/grp<br>eyecup<br>lysates |
| 2 | CTx001 Low | 9 rats/18<br>eyes | Yes | 1x10<sup>8</sup>vg/eye,<br>subretinal | 2.5 µL | At day 0<br>and day 1 |  |
| 3 | CTx001 High | 9 rats/18<br>eyes | Yes | 5x10<sup>9</sup>vg/eye,<br>subretinal | 2.5 µL | At day 0<br>and day 1 |  |

**Supplementary Figure 1.**
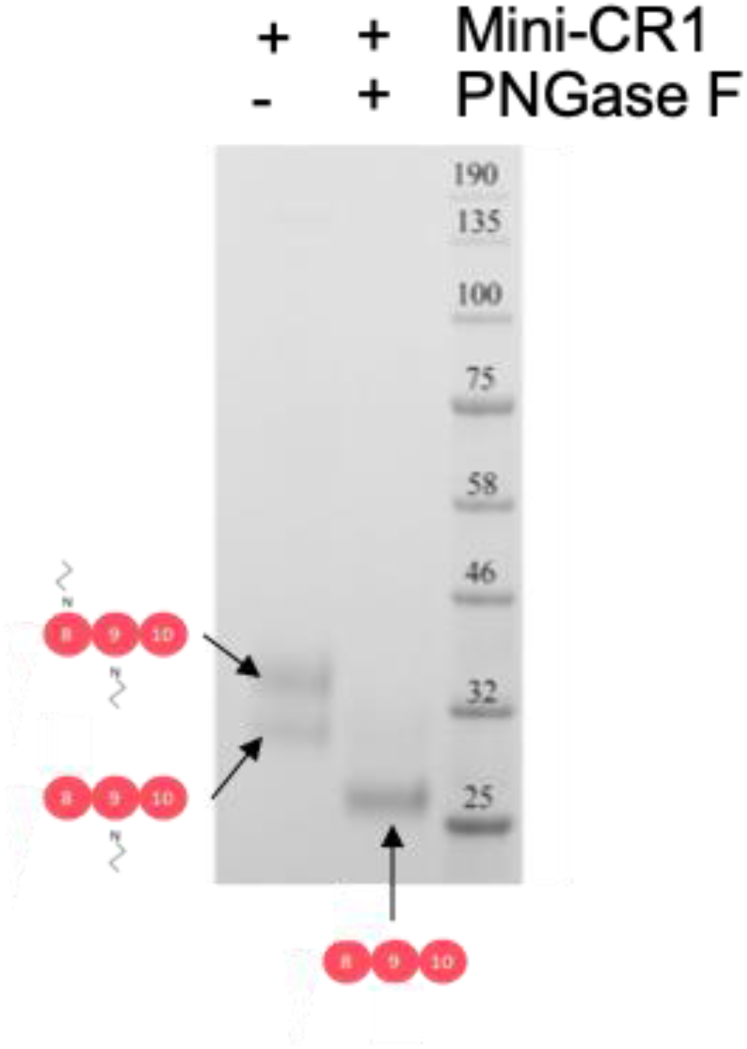
Glycosylation of mini-CR1 and its ability to generate C3dg. A) Purified recombinant mini-CR1 runs as two separate bands on a reducing SDSPAGE 4-12% gel and visualised with Coomassie Blue stain. Both bands resolve into a single band after removal of N-linked glycosylation by treatment with PNGaseF. Data shown representative of three independent experiments.

**Supplementary Figure 2.**
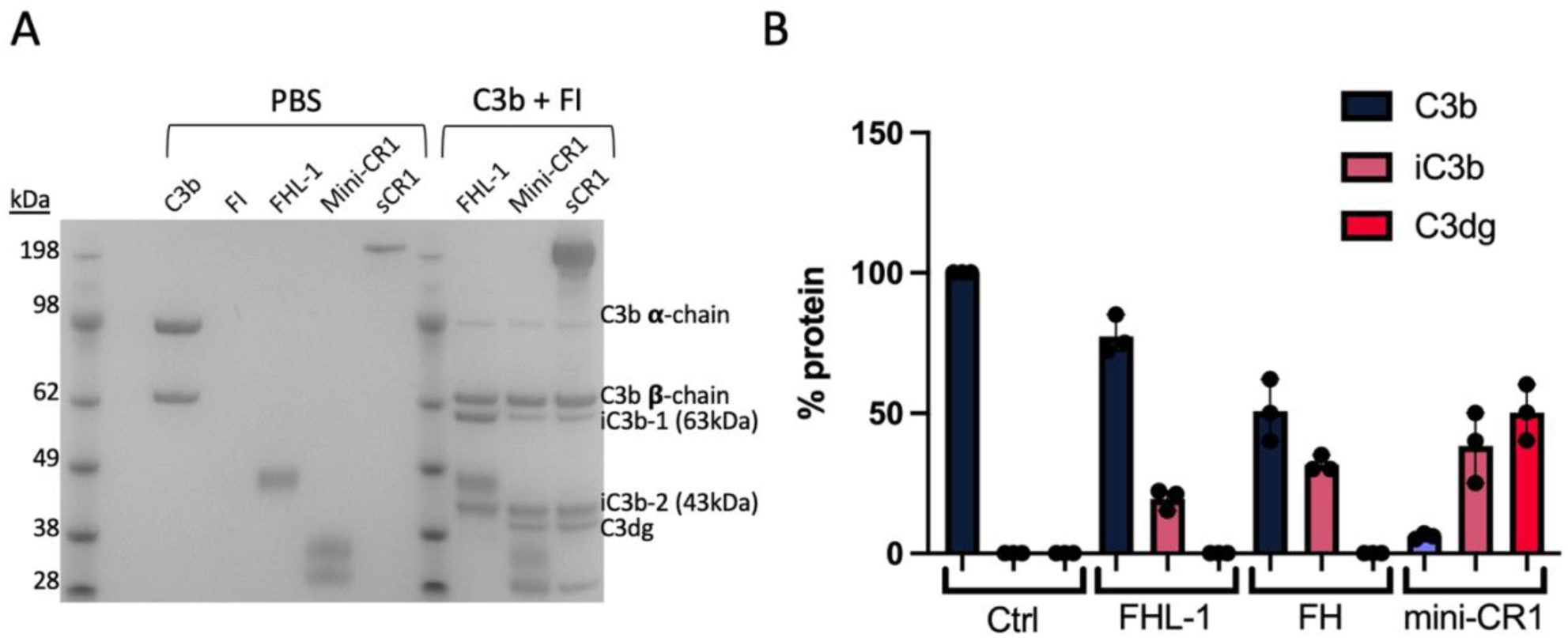
Fluid phase C3b cleavage assays in the presence of FI and different cofactors. A) SDS-PAGE with Coomassie blue staining showing effects of equimolar concentrations (1µM) of FHL-1, mini-CR1 and soluble CR1 (sCR1) mixed with 2µg C3b and 0.04µg FI. The presence of mini-CR1 or sCR1 resulted in C3dg formation whereas FHL-1 did not. B) Band densitometry was performed on Coomassie blue stained gels and the conversion of C3b into iC3b and C3dg is reported as total percentage protein (i.e. compared to band density of C3b alone); Control (Ctrl) FI alone, and with FHL-1, FH and mini-CR1 added.

**Supplementary Figure 3.**
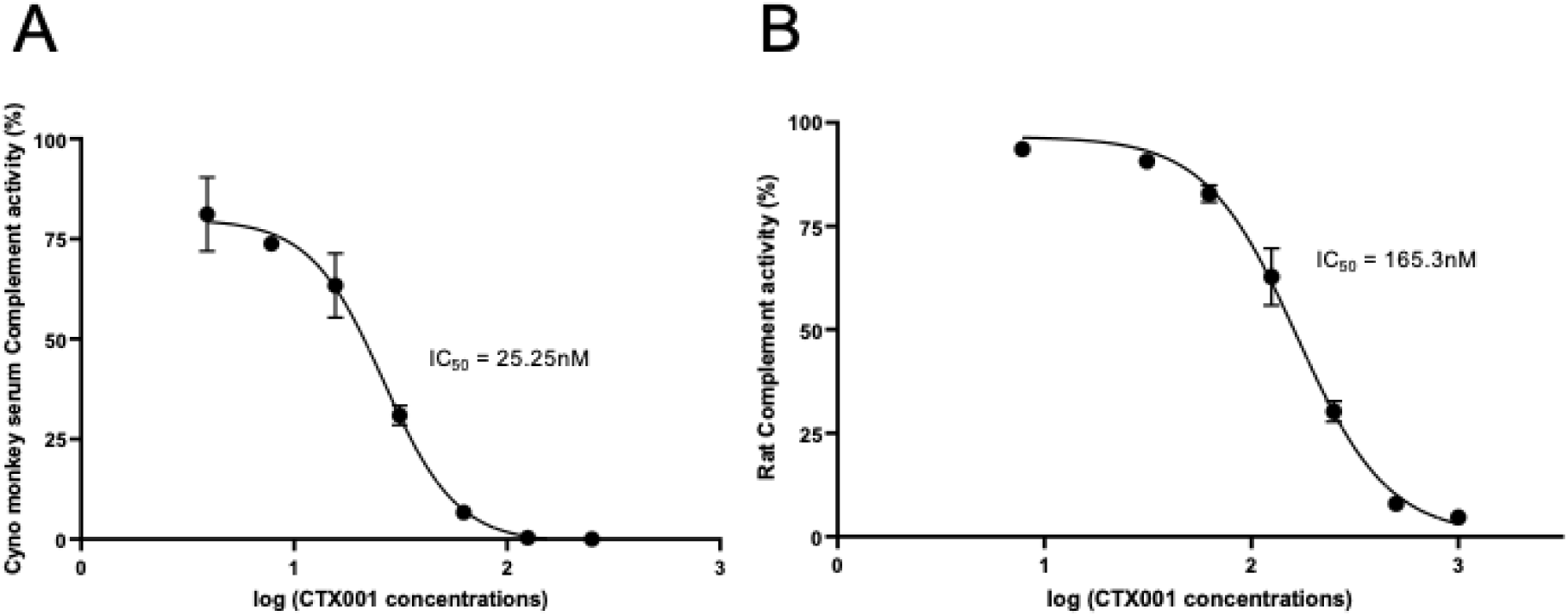
IC50 analysis for mini-CR1 in NHP and rat serum. By way of comparison, purified recombinant mini-CR1 was tested for efficacy in inhibiting the alternative pathway of complement in the serum of NHP (A) or rat (B)using commercially available Weisslab assays: representative graphs from three independent experiments are shown.

**Supplemental Figure 4.**
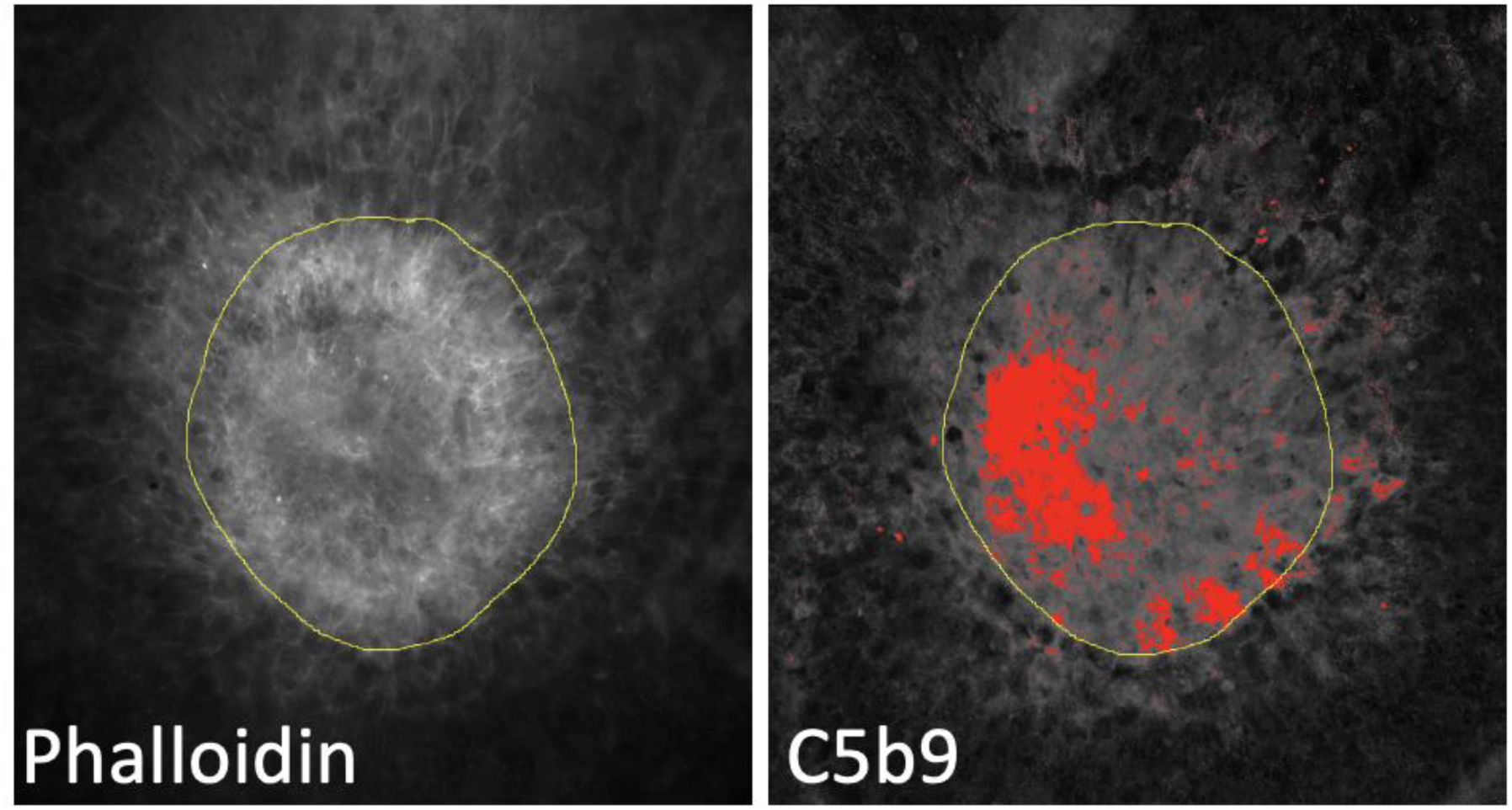
Quantitative image analysis of C5b9 staining within the laser lesion ROI.

## Notes

**Sources of support for work:** This work was supported by an MRC CDA Fellowship (MR/K024418/1) and Complement Therapeutics.

## References

1. Colijn, J. M. et al. Prevalence of Age-Related Macular Degeneration in Europe: The Past and the Future. Ophthalmology 124, 1753–1763 (2017).

2. Wong, W. L. et al. Global prevalence of age-related macular degeneration and disease burden projection for 2020 and 2040: A systematic review and meta-analysis. Lancet Glob Health 2, e106–16 (2014).

3. Hristodorov, D., Lohoff, T., Luneborg, N., Mulder, G.-J. & Clark, S. J. Investing in vision: Innovation in retinal therapeutics and the influence on venture capital investment. Prog Retin Eye Res 99, 101243 (2024).

4. Rathi, S., Hasan, R., Ueffing, M. & Clark, S. J. Therapeutic targeting of the complement system in ocular disease. Drug Discov Today 28, 103757 (2023).

5. Whitmore, S. S. et al. Complement activation and choriocapillaris loss in early AMD: Implications for pathophysiology and therapy. Prog Retin Eye Res 45, 1–29 (2015).

6. Mullins, R. F. et al. The Membrane Attack Complex in Aging Human Choriocapillaris. Am J Pathol 184, 3142–3153 (2014).

7. Mullins, R. F. et al. Elevated membrane attack complex in human choroid with high risk complement factor H genotypes. Exp Eye Res 93, 565–567 (2011).

8. Sohn, E. H. et al. Choriocapillaris Degeneration in Geographic Atrophy. American Journal of Pathology 189, 1473–1480 (2019).

9. Cipriani, V. et al. Beyond Factor H: the influence of genetic variation with age-related macular degeneration on circulating Factor H-like 1 and Factor H-related protein levels. Am J Hum Genet 108, 1385–1400 (2021).

10. Cipriani, V. et al. Increased circulating levels of Factor H-Related Protein 4 are strongly associated with age-related macular degeneration. Nat Commun 11, 778 (2020).

11. Lorés-Motta, L., et al. Common haplotypes at the CFH locus and low-frequency variants in CFHR2 and CFHR5 associate with systemic FHR concentrations and age-related macular degeneration. The American Journal of Human Genetics 108, 1367–1384 (2021).

12. Reeve, M. P. et al. Loss of CFHR5 function reduces the risk for age-related macular degeneration. medRxiv 2024.11.11.24317117 (2024).

13. Emilsson, V. et al. A proteogenomic signature of age-related macular degeneration in blood. Nat Commun 13, 3401 (2022).

14. Choudhury, R. et al. FHL-1 interacts with human RPE cells through the α5β1 integrin and confers protection against oxidative stress. Sci Rep 11, 1–17 (2021).

15. Clark, S. J. et al. His-384 allotypic variant of factor H associated with age-related macular degeneration has different heparin binding properties from the non-disease-associated form. Journal of Biological Chemistry 281, 24713–24720 (2006).

16. Clark, S. J., McHarg, S., Tilakaratna, V., Brace, N. & Bishop, P. N. Bruch’s membrane compartmentalizes complement regulation in the eye with implications for therapeutic design in age-related macular degeneration. Front Immunol 8, 1778 (2017).

17. Seya, T. & Atkinson, J. P. Functional properties of membrane cofactor protein of complement. Biochemical Journal 264, 581–588 (1989).

18. Forneris, F. et al. Regulators of complement activity mediate inhibitory mechanisms through a common C3b-binding mode. EMBO J 35, 1133–1149 (2016).

19. Xue, X. et al. Regulator-dependent mechanisms of C3b processing by factor i allow differentiation of immune responses. Nat Struct Mol Biol 24, 643–651 (2017).

20. Lambris, J. D. et al. Dissection of CR1, factor H, membrane cofactor protein, and factor B binding and functional sites in the third complement component. The Journal of Immunology 156, 4821–4832 (1996).

21. McHarg, S., Brace, N., Bishop, P. N. & Clark, S. J. Enrichment of Bruch’s Membrane from Human Donor Eyes. Journal of Visualized Experiments e53382 (2015).

22. Kreideweiss, S. et al. Pharmacological characterization of the novel complement C3 inhibitor BI-C. Invest Ophthalmol Vis Sci 65, 1952–1952 (2024).

23. Vandenberghe, L. H. et al. Dosage thresholds for AAV2 and AAV8 photoreceptor gene therapy in monkey. Sci Transl Med 3, 88ra54 (2011).

24. Fearon, D. T. & Austen, K. F. Activation of the alternative complement pathway due to resistance of zymosan bound amplification convertase to endogenous regulatory mechanisms. Proc Natl Acad Sci U S A 74, 1683–1687 (1977).

25. Morrison, D. C., Kline, L. F. & Lideritz, O. Activation of the Classical and Properdin Pathways of Complement by Bacterial Lipopolysaccharides (LPS). The Journal of Immunology 118, 362–368 (1977).

26. Tosic, L., Sutherland, W. M., Kurek, J., Edberg, J. C. & Taylor, R. P. Preparation of monoclonal antibodies to C3b by immunization with C3b(i)-Sepharose. J Immunol Methods 120, 241–249 (1989).

27. Bayly-Jones, C. et al. The neoepitope of the complement C5b-9 Membrane Attack Complex is formed by proximity of adjacent ancillary regions of C9. Commun Biol 6, 42 (2023).

28. Tobe, T. et al. Targeted disruption of the FGF2 gene does not prevent choroidal neovascularization in a murine model. American Journal of Pathology 153, 1641–1646 (1998).

29. Kwak, N., Okamoto, N., Wood, J. M. & Campochiaro, P. A. VEGF is major stimulator in model of choroidal neovascularization. Invest Ophthalmol Vis Sci 41, 3158–3164 (2000).

30. Liu, J. et al. Relationship between complement membrane attack complex, chemokine (C-C motif) ligand 2 (CCL2) and vascular endothelial growth factor in mouse model of laser-induced choroidal neovascularization. Journal of Biological Chemistry 286, 20991–21001 (2011).

31. Bora, P. S. et al. Role of Complement and Complement Membrane Attack Complex in Laser-Induced Choroidal Neovascularization. The Journal of Immunology 174, 491– 497 (2005).

32. Cashman, S. M., Ramo, K. & Kumar-Singh, R. A Non Membrane-Targeted Human Soluble CD59 Attenuates Choroidal Neovascularization in a Model of Age Related Macular Degeneration. PLoS One 6, e19078 (2011).

33. Lyzogubov, V. V. et al. Role of Ocular Complement Factor H in a Murine Model of Choroidal Neovascularization. Am J Pathol 177, 1870–1880 (2010).

34. Stampoulis, D. et al. In vivo expression and efficacy following subretinal delivery of GT005, an investigational gene therapy for the treatment of geographic atrophy (GA). Invest Ophthalmol Vis Sci 63, 293 (2022).

35. Lachmann, P. J. Preparing serum for functional complement assays. J Immunol Methods 352, 195–197 (2010).

36. Wildner, G. Are rats more human than mice? Immunobiology 224, 172–176 (2019).

37. Dalkara, D. et al. In vivo-directed evolution of a new adeno-associated virus for therapeutic outer retinal gene delivery from the vitreous. Sci Transl Med 5, 189ra76 (2013).

38. Nakaizumi, Y. The Ultrastructure of Bruch’s Membrane: I. Human, Monkey, Rabbit, Guinea Pig, and Rat Eyes. Archives of Ophthalmology 72, 380–387 (1964).

39. Csaky, K. et al. New approaches to the treatment of Age-Related Macular Degeneration (AMD). Exp Eye Res 221, 109134 (2022).

40. Hammadi, S. et al. Bruch’s Membrane: A Key Consideration with Complement-Based Therapies for Age-Related Macular Degeneration. J Clin Med 12, 2870 (2023).

41. Clark, S. J. et al. Breaking Bruch’s: How changes in Bruch’s membrane influence retinal homeostasis. Exp Eye Res 255, 110343 (2025).

42. Lucientes-Continente, L., Márquez-Tirado, B. & Goicoechea de Jorge, E. The Factor H protein family: The switchers of the complement alternative pathway. Immunol Rev 313, 25–45 (2023).

43. Kárpáti, É. et al. Interaction of the Factor H Family Proteins FHR-1 and FHR-5 With DNA and Dead Cells: Implications for the Regulation of Complement Activation and Opsonization. Front Immunol 11, 1297 (2020).

44. Zouache, M. A. et al. Levels of complement factor H-related 4 protein do not influence susceptibility to age-related macular degeneration or its course of progression. Nat Commun 15, 1–17 (2024).

45. Morgan, P CD59. in The Complement FactsBook: Second Edition 361–367 (Academic Press, 2018). doi:10.1016/B978-0-12-810420-0.00034-1.

46. de Jong, S., Tang, J. & Clark, S. J. Age-related macular degeneration: A disease of extracellular complement amplification. Immunol Rev 313, 279–297 (2023).

